# Automated cell type annotation and exploration of single cell signalling dynamics using mass cytometry

**DOI:** 10.1101/2022.08.13.503587

**Authors:** Dimitrios Kleftogiannnis, Sonia Gavasso, Benedicte Sjo Tislevoll, Nisha van der Meer, Inga K. F. Motzfeldt, Monica Hellesøy, Stein-Erik Gullaksen, Emmanuel Griessinger, Oda Fagerholt, Andrea Lenartova, Yngvar Fløisand, Bjørn Tore Gjertsen, Inge Jonassen

## Abstract

Mass cytometry by time-of-flight (CyTOF) is an emerging technology allowing for in-depth characterisation of cellular heterogeneity in cancer and other diseases. However, computational identification of cell populations from CyTOF, and utilisation of single cell data for biomarker discoveries faces several technical limitations, and although some computational approaches are available, high-dimensional analyses of single cell data remains quite demanding. Here, we deploy a bioinformatics framework that tackles two fundamental problems in CyTOF analyses namely: a) automated annotation of cell populations guided by a reference dataset, and b) systematic utilisation of single cell data for more effective patient stratification. By applying this framework on several publicly available datasets, we demonstrate that the Scaffold approach achieves good tradeoff between sensitivity and specificity for automated cell type annotation. Additionally, a case study focusing on a cohort of 43 leukemia patients, reported salient interactions between signalling proteins that are sufficient to predict short-term survival at time of diagnosis using the XGBoost algorithm. Our work introduces an automated and versatile analysis framework for CyTOF data with many applications in future precision medicine projects. Datasets and codes are publicly available at: https://github.com/dkleftogi/singleCellClassification

## Introduction

Mass cytometry by time-of-flight (CyTOF) is a powerful technology that allows for simultaneous quantification of over 30 parameters/proteins with single cell resolution (1–2). The application of CyTOF has revolutionised our understanding about cellular biology and immunology, and several studies have emphasized on the role of cellular phenotypic heterogeneity in human diseases (3–4).

Although the CyTOF data acquisition is conceptually simple, the analysis of high-dimensional single cell data faces several technical limitations. Typically, CyTOF downstream analysis frameworks require accurate identification of cellular populations (i.e., equally denoted as phenotypes or cell types). For this task ‘manual gating’ based on bi-axial plots is the simplest approach borrowed from the conventional flow cytometry field (5). However, in CyTOF analyses due to the high-dimensional data, manual gating is impractical and tedious (6). To mitigate this limitation, several unsupervised clustering algorithms have been deployed aiming at partitioning cells to clusters and assigning clusters to biological cell populations (7–10). Cell type annotation via clustering, although widely applicable, faces also technical challenges. Clustering itself is an ill-defined computational problem and algorithms are sensitive to their internal parameter which makes them difficult to tune. In addition, annotation of clusters of cells that correspond to biological cell populations is performed post hoc, leading to a labour-intensive process that is subjective to decisions from the researcher/analyst. In the era of ‘big data’, the availability of ‘well-annotated’ datasets has opened unprecedented opportunities for automated cell type annotation via Machine Learning (ML) algorithms (11–13). For example, it has been shown that supervised ML algorithms trained with single cells from annotated cell types, achieve high precision (6), whereas semi-supervised approaches (14–15) gain popularity due to their ability to delineate the structure of the data and identify ‘rare’ cell types. Consequently, exploiting annotated datasets using projections to reference datasets (referred hereafter as reference-guided annotation) or other similar approaches that ‘align’ cells to reference datasets is a promising research direction for visualisation and automated cell type annotation for large scale data exploration (16–20).

Subsequently, once the cell type annotation step has been completed, CyTOF data can be used to minutely analyse biological cell populations and answer biologically and clinically relevant questions. For example, both in cancer and many other diseases, predicting responders from non-responders to therapies might improve the quality of patients’ life. Hence, many precision medicine studies have been presented combining CyTOF data, clinical information, and/or genetic data with Machine Learning (ML) algorithms (21–24).

Using CyTOF data, detecting changes in phospho-signalling protein expression levels, and dissecting their dynamics holds a great promising towards the implementation of precision medicine. Term ‘dynamics’ reflects changes in the expression levels between proteins because of the disease condition. However, engineering phospho-signalling features with conventional statistical approaches such as mean, median or quantile expression values neglect rich information measured at the single cell level. To overcome this limitation, approaches such as the Statistical Analysis of Perturbation Responses (SARA) (7), or the Density Resampled Estimate of Mutual Information (DREMI) have been proposed (25). However, to the best of our knowledge, the latter approaches have not been fully exploited with ML algorithms to identify cell type-specific signalling dynamics that are predictive of patients’ survival. To note.

Here we present a novel bioinformatics framework that tackles two fundamental problems in CyTOF analyses namely: a) automated cell type annotation guided using reference datasets; and b) systematic utilisation of protein dynamics measured with CyTOF for patient stratification. Our experimentation with several publicly available datasets and ML algorithms shows that the Scaffold map approach combined with a comprehensive proteo-genomic reference map of the human blood and bone marrow (26), achieves superior performance compared to other supervised and semi-supervised ML learning approaches. With the developed framework we also show that cell type-specific DREMI scores combined with the XGBoost algorithm can be used to discriminate accurately long-term from short-term survivors in leukemia. We anticipate that the proposed analysis framework could serve as a paradigm for future personalised medicine projects.

## Results

### Reference map for automated cell type annotation and self-consistency test

Previous studies have shown that supervised and semi-supervised ML methods such as the Linear Discriminant Analysis (LDA), the K-nearest neighbour (KNN) or CyAnno are highly effective for cell type annotation. These approaches require a well-annotated set of single cells to be used for training, but alike all other ML-based approaches they often perform poorly in testing data that are totally unseen from the training data. In a seminal study Spitzer et al. presented the Scaffold approach that was used for visualisation and automated cell type annotation (27). The Scaffold approach builds a reference map from single cells, and it is quite flexible allowing mapping of new datasets that can be even acquired from different modalities/technologies. With this idea in mind, here we combine the Scaffold approach with a comprehensive multimodal reference dataset acquired from Abseq experiments measuring simultaneously mRNA and surface protein expression in single cells (26). Supplementary Figure 1 provides more information about the reference dataset including the original dimensionality reduction UMAP plot (e.g., annotation and marker expression overlayed), the relative abundance of the available cell types, as well as a graphical representation of the force-directed graph generated using the Scaffold.

First, to assess the relative performance of different ML-based cell type annotation methods, we perform a self-consistency set focusing on the reference dataset. We utilise 80% of the available cells for training, we keep the remaining 20% of the available cells for testing, and we estimate Accuracy, Sensitivity, Specificity and F1 in a multi-class setting for the available cell types (formulas available in Methods). Figure 1a shows that the Scaffold map approach achieves perfect classification followed by the CyAnno approach that is overall ranked second. Two versions of KNN with K=3 and K=10 as well as LDA achieve the lowest classification performance, and the results are consistent in an additional experiment where 50% of the available cells are used for training and the remaining 50% of the available cells are kept aside for testing (Supplementary Figure 2).

**Figure 1:**
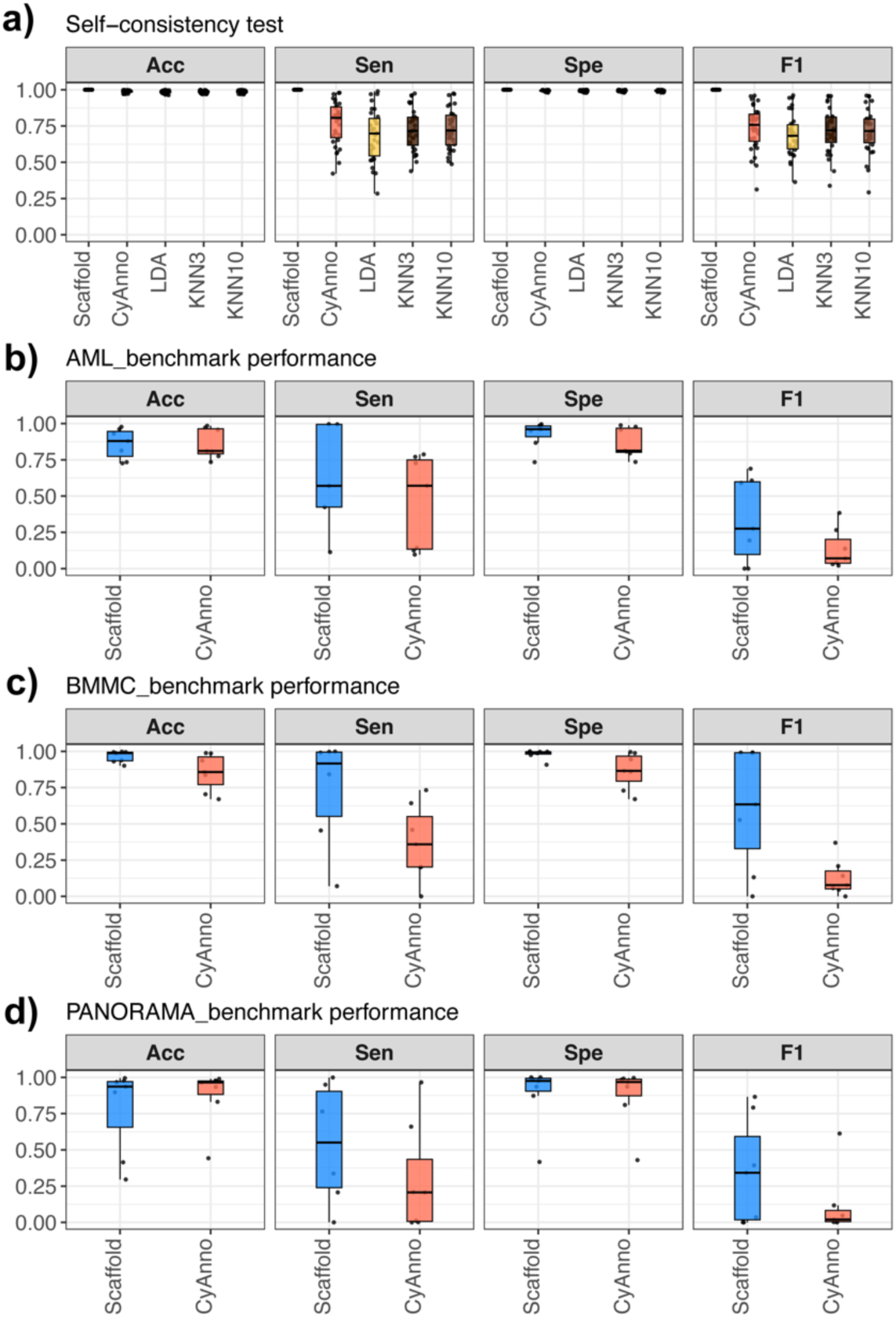
Performance evaluation of different ML-based methods for automated cell type annotation. a) Results summarising the classification performance of Scaffold, CyAnno, LDA and KNN with K=3 and KNN with K=10, across 31 cell types from the Reference dataset; b-d) Results summarising the classification performance of Scaffold and CyAnno for the prediction of seven cell types from: AML_benchmark dataset (b), BMMC_benchmark dataset (c) and PANORAMA_benchmark dataset(d). The performance is assessed using Accuracy (ACC), Sensitivity (Sen), Specificity (Spe) and F1 score.

### Assessing the performance of the Scaffold approach using independent mass cytometry datasets

Next, to assess the performance of the Scaffold approach we design a more realistic experimental scenario. We downloaded three manually gated CyTOF datasets, that are considered benchmarks for similar studies (6,11). Supplementary Figure 3 and Supplementary Figure 4 show details about the benchmark datasets, denoted here as AML_benchmark, BMMC_benchmark and PANORAMA_benchmark. During this experimentation the full reference dataset from Triana et al. is used as ‘training’ set, and the cell type classification performance is measured in the benchmark datasets. This experimentation provides a completely unbiased way of assessing the methods’ effectiveness since the testing sets are completely independent and unseen from the reference dataset.

Similar to the previous subsection Accuracy, Sensitivity, Specificity and F1 are computed in a multi-class setting across the available cell types for CyAnno and the Scaffold approaches since they are top-2 performers of the self-consistency test. Figure 1b-d shows that the Scaffold approach surpasses CyAnno in all performance metrics. By inspecting more closely the outcome for both methods we also observe that for some cell types the performance is rather low. Considering the F1 score as a representative performance metric we find that both methods fail to detect NK_CD16pos cells in the AML_benchmark, pDCs and HSCs_MPPs cells in the BMMC_benchmark and pDCs and NK_CD16pos cells in the PANORAMA_benchmark. One possible explanation could be attributed to the lack of more appropriate antibodies in the benchmark datasets to be able to detect accurately these cell types. To control for this limitation, it is highly advisable for users to inspect the ‘automated’ predictions in totally unseen datasets and consider the expression profiles of cell-type defining markers to eliminate false positives and false negative predictions. Taken together the results from this analysis demonstrate that the Scaffold approach built on top of a comprehensive reference dataset achieves in most cases good trade-off between sensitivity and specificity for cell types, where the reference dataset (e.g., training set) is completely independent from the testing data. Consequently, it opens possibilities for automated and scalable data-driven exploration of clinical CyTOF dataset that we will show in the next sections.

### Precision medicine application of the developed framework

To showcase the use of the developed bioinformatics framework in a precision oncology application we perform a case study using publicly available CyTOF dataset from leukemia (21). To increase cohort size, the published dataset is toped-up with additional leukemia samples that were collected and processed together with the published cohort from ref. 21. This results to 43 samples from leukemia patients collected at time of diagnosis that are accompanied with clinical and genetic information. For comparison purposes we also downloaded and analysed a cohort of seven healthy donors that was used as a control in the original publication (21).

### 3.1 Phenotyping data from leukemia patients and healthy donors

First, the Scaffold approach is used to annotate the cell populations guided by the reference dataset from Triana et al. (26). Given the common antibodies between the panels of the two datasets (the following common markers are considered ‘backbone’: CD11b, CD8a, CD33, CD34, CD3, CD123, CD56, CD14, CD117, CD38, CD4, CD16 C D20, CD45, CD7) we can detect seven cell populations namely: B cells, CD4+ T cells (CD4_T), CD8+ T cells (CD8_T), Hematopoietic Stem Cells and Multipotent Progenitor cells (HSCs_MPPs), Monocytes, Natural Killer cells (NK) and plasmocytoid Dendritic Cells (pDCs). Supplementary Figure 5 shows a UMAP representation of selected cell populations from the reference dataset, as well as UMAP projections of leukemia cells and healthy control cells to the reference UMAP using the Scaffold approach. Clear separation of the cell types is observed in all cases, whereas in the leukemia case an expansion of the HSCs_MPPs cell population compared to the healthy controls is visually detected. This cell population probably represents leukemic ‘blasts’ that are immature white blood cells with low expression for CD45 and high expression of CD34 markers. For confirmation, Supplementary Figure 6-8 shows the expression levels of important cell type-defining markers for all datasets, and consistent expression profiles based on the literature are detected.

Once all cells from patients and healthy controls are annotated, we compare their relative cellular abundances using statistical approaches (Figure 2a). Statistically different distributions are found for CD8+ T cells (CD8_T), the Hematopoietic Stem Cells and Multipotent Progenitor cells (HSCs_MPPs), and the Natural Killer (NK) cells. As seen before, the HSCs_MPPs cell type in leukemia is expanded compared to healthy controls, whereas the abundance of CD8+ T and NK cells is lower in leukemia. As expected, the cellular abundance distributions in leukemia appear quite heterogeneous compared to healthy controls.

**Figure 2:**
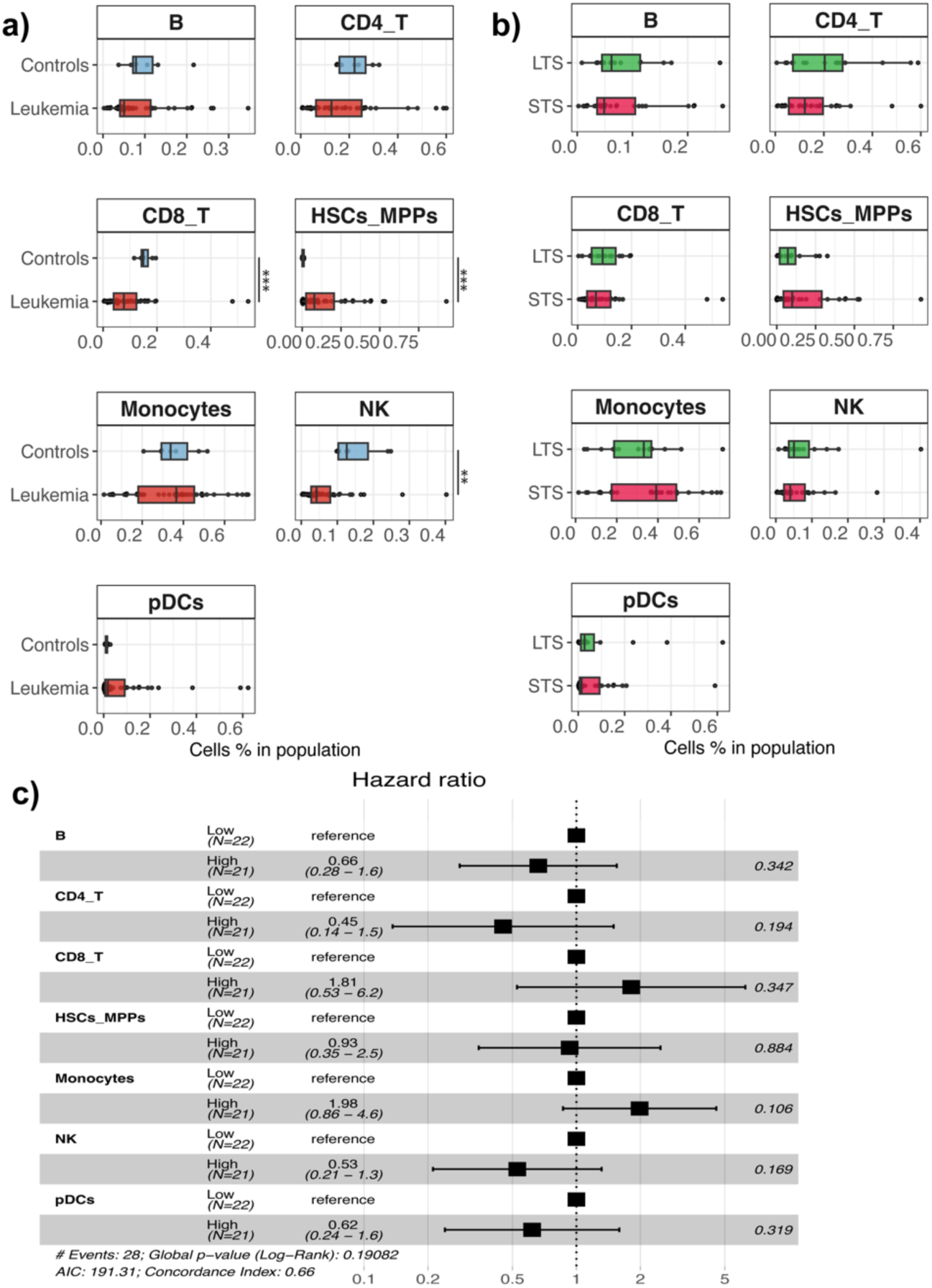
Comparing cellular abundance distributions between leukemia patients and healthy controls. a) The boxplots show the cellular abundance distributions of 43 patients and seven healthy donors; b) The boxplots show the cellular abundance distributions of patients with short-term survival (n=28, STS) and long-term survival (n=15, LTS). c) Cox proportional hazards modelling using as input the cellular abundances (%) measured at time of diagnosis. Statistically significant differences were tested using Wilcoxon rank sum test. The levels of significance are: * for p<0.05, ** for p<0.01 and *** for p<0.001.

Next, we focus on the available leukemia samples by incorporating clinical information for stratification. We utilise 5-year survival information to stratify patients, into two groups (that we hereafter refer to as “Long Term Survivors” (LTS, n=15), with survival equal/greater than 60 months after diagnosis, and “Short Term Survivors” (STS, n=28) with survival shorter than 60 months after diagnosis. More details about the demographics of this dataset can be found in ref. (21) and in Supplementary Table 1 and Supplementary Table 2. We compare the cellular abundance distributions between STS and LTS groups (Figure 2b), but we are not able to detect statistically significant differences. To investigate further possible associations with patients’ survival, the relative cellular abundances of STS and LTS groups are discretised based on the median cohort value and used as input for multivariate Cox proportional hazard modelling. Again, no statistically significant differences are detected between the groups (Figure 2c, Schoenfeld residuals are shown in Supplementary Figure 9). Finally, to explore possible associations between genetic information and patients’ cellular abundances we incorporate the karyotype information and the FLT3-ITD mutational status which is a well-known hotspot mutation in leukemia. Supplementary Figure 10 summarises the results. Considering the karyotype, we are not able to detect statistically significant differences between the groups after adjusting the p-values for multiple comparisons. When considering the FLT3-ITD status (wild type vs. mutated) we find that the mutated patients have significantly lower abundance of CD8+ T cells (CD8_T) and CD4+ T (CD4_T) cells, whereas they have significantly higher abundance of Natural Killer (NK) and plasmacytoid Dendritic cells (pDCs). However, due to the small cohort size the statistical power is limited. Thus, the results must be generalised with caution, as after stratification and comparisons between groups, some distributions included less than 10 data points.

In summary, the deployed cell type annotation framework, allows automated comparisons of cellular abundances between patient subgroups enabling incorporation of clinical and genetic information for patient stratification.

### 3.2 Predicting patients’ survival with machine learning

Predicting patients’ survival using single cell omics datasets is one of the biggest challenges in modern biomedical research. So far, we have utilised relative cellular abundances to compare STS from LTS patients, but in most of the cases our results did not reach statistically significant levels. In this subsection we hypothesize that utilising more advanced ML algorithms applied on features extracted from the ‘rich’ single cell data may lead to improved patient stratification and more effective survival prediction. To explore this hypothesis, we design a ML-based bioinformatics framework that is built on top of the automated cell type annotation step. Supplementary Figure 11 provides a graphical abstract of the ML-based framework and the performance evaluation setup using Sensitivity, Specificity, F1 and Area Under the Curve (AUC) metrics (formulas described in Methods).

#### 3.2.1 Assessing the performance of median expression values as features for machine learning

First, we utilise the signalling markers of the antibody panel to generate features using the median expression value for each marker. With this formulation the problem of discriminating STS from LTS is a binary classification task and different algorithms are tested including XGBoost (28), LASSO and Ridge regression, using as input the constructed feature matrices (29). ML-based models are trained separately for every of the available cell types, and the classification performance is recorded in the test set (20% of instances) that is completely independent from the training/validation set (80% of instances). To ensure reproducibility, the whole training/validation and testing was repeated 100 times with random split between the sets.

An important problem related to most biomedical datasets is the imbalance of positive and negative classes. Specifically for the prediction of survival in cancer research, the class imbalance problem is present as one of the classes (e.g., STS) usually outnumbers the other (e.g., LTS). To mitigate this problem over/under sampling techniques have been proposed with Synthetic Minority Over-sampling TEchnique (SMOTE) being one of the most effective (30). To deal with the class imbalance problem in our case study we apply SMOTE, and the training and testing procedures are repeated with different ratios between STS and LTS data instances. Figure 3 summarises the results of our experimentation.

**Figure 3:**
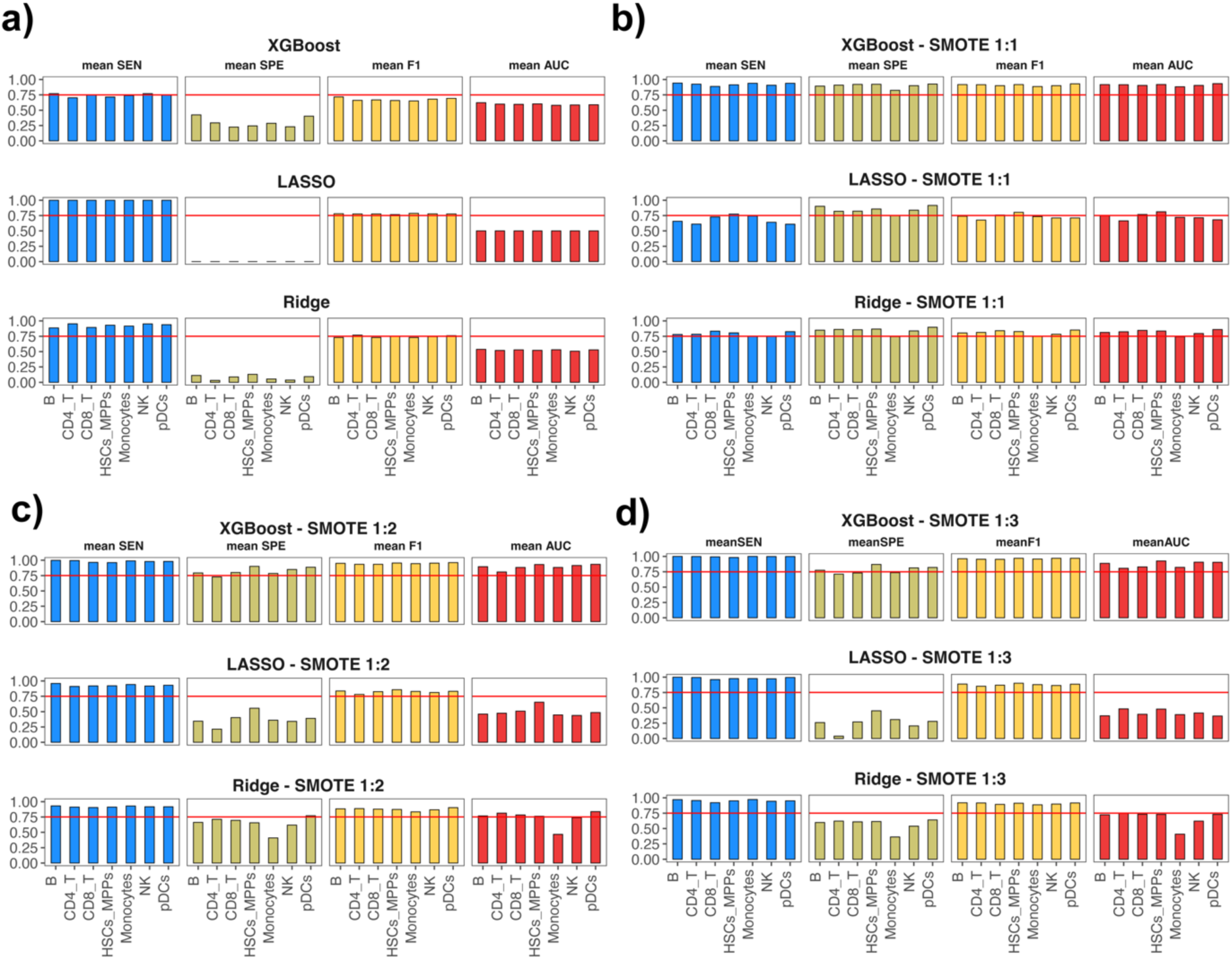
Performance evaluation of different ML-based methods for survival prediction in leukemia using as features median expression values of phospho-proteins. Average classification performance of XGBoost, LASSO and Ridge regression for discriminating STS from LTS patients using data from: a) the original case study cohort; b) SMOTE synthetic data with ratio 1:1 between the classes; c) SMOTE synthetic data with ratio 1:2 between the classes; and d) SMOTE synthetic data with ratio 1:3 between the classes. The performance is assessed using mean Sensitivity (SEN), mean Specificity (SPE), mean F1 and mean Area Under Curve (AUC) of 100 executions with random splits between training/validation and testing sets.

Our results show that it is possible to discriminate STS from LTS with high levels of success using cell type-specific ML models and median expression values of signalling proteins in the CyTOF panel as input features. We also demonstrate that the class imbalance problem affects drastically the predictive performance of the developed ML models. Within our case study, the SMOTE ratio 1:1 achieves the highest geometric mean of sensitivity and specificity for all classification algorithms tested. Focusing on different classification algorithms we find that XGBoost surpasses the regularised regression methods LASSO and Ridge and achieves the best performance that reaches an average Sensitivity of 0.92, average Specificity of 0.89, average F1 of 0.90 and average AUC of 0.91. Finally, our findings are robust when the whole experimentation is repeated with 50-50 split between training/validation and testing subsets (Supplementary Figure 12).

#### 3.2.2 Using DREMI scores to engineer features for machine learing-based modelling of survival

In this subsection we rationalise that utilising Conditional Density Resampled Estimate of Mutual Information known as DREMI (25), to engineer features for ML-based prediction of survival may lead to superior classification performance.

As a proof of principle, we use DREMI scores to engineer features and developed cell type-specific models to predict patients’ survival. DREMI operates on pairs of signalling proteins and quantifies the ‘strength’ of the relationship between these two proteins meaning that it provides a numerical score to assess how the expression levels of protein X affects the expression levels of protein Y in a signalling cascade where protein X interacts with Y (denoted as X→Y). An inherited drawback of this approach is the directionality of the relationships, meaning that the DREMI score between proteins X → Y is in principle different from the score Y → X. To alleviate this limitation and develop a more general bioinformatics framework we quantify all possible pairs between proteins in the antibody panel. This results to engineering 210 features using all possible combinations of 15 available signalling proteins in the panel counting both directions.

The experimentation presented in the previous subsection is repeated using the XGBoost algorithm, deploying SMOTE to account for the effect of class imbalance in the datasets. Our results show (Figure 4a) that the combination of DREMI features with the XGBoost algorithm, achieves higher classification performance compared to the median-derived features trained and tested with the same algorithm. In addition, our experimentation with SMOTE reconfirms that the class imbalance impacts the classification performance of the developed ML models. Within our experimentation the SMOTE ratio 1:1 achieves the highest performance across the available cell types that reaches an average Sensitivity of 0.98, average Specificity of 0.97, average F1 of 0.98 and average AUC of 0.98.

**Figure 4:**
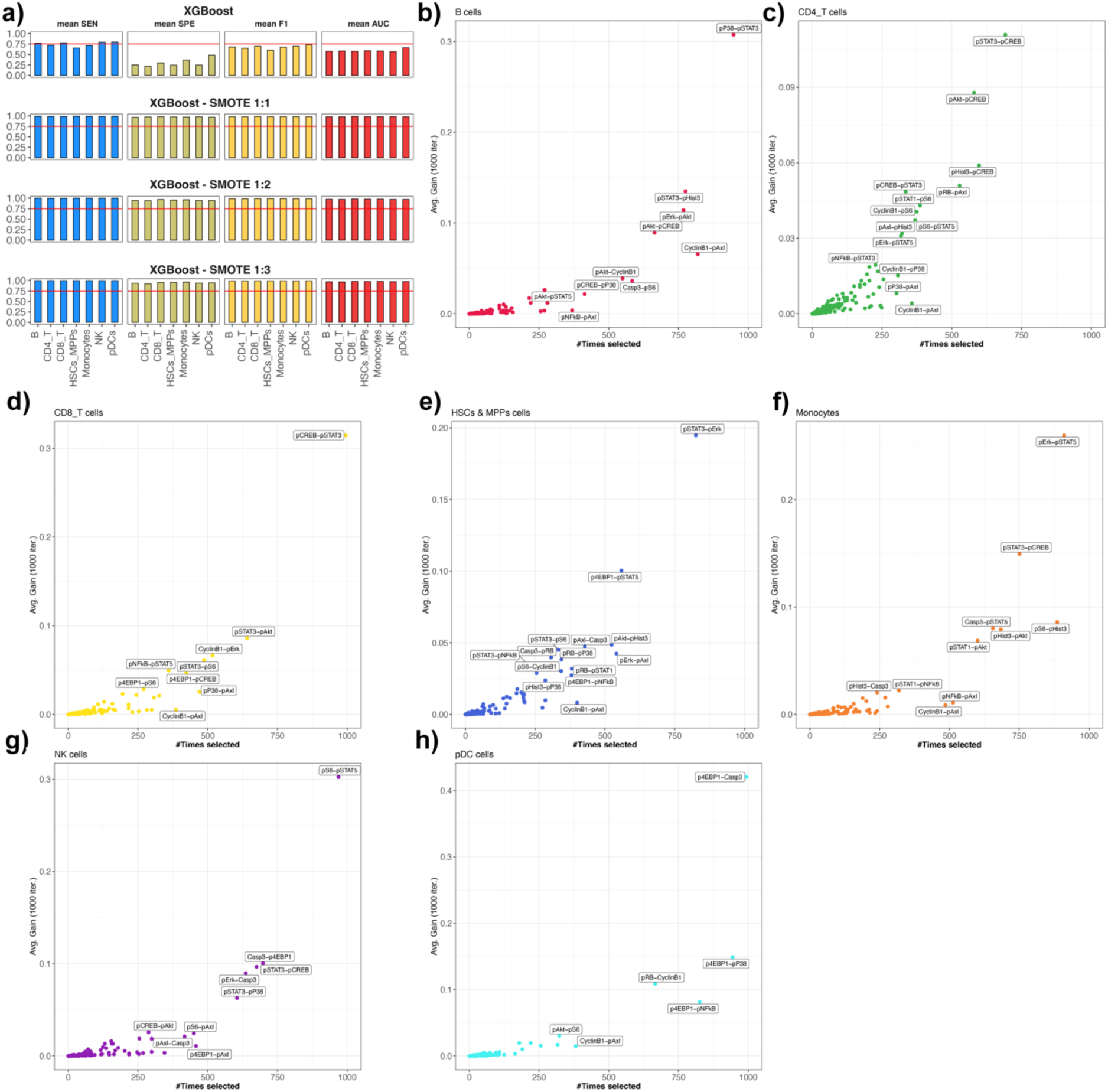
Performance evaluation and feature importance using DREMI-derived features. a) Average classification performance of XGBoost algorithm for discriminating STS from LTS patients trained/validated and tested using the original data as well as SMOTE-derived datasets with different ratios between the class. The performance is assessed using mean Sensitivity (SEN), mean Specificity (SPE), mean F1 and mean Area Under Curve (AUC) of 1000 executions with random splits between training/validation and testing sets; b-h) Scatter plots showing the average feature gain as returned by the XGBoost algorithm (Y axis) and the number of times each feature is selected in independent execution runs (X axis). Each subplot corresponds to cell type-specific development of ML models.

#### 3.2.3 Studying the feature importance from learnt models

One of the biggest advantages of the XGBoost algorithm is its’ ability to retrieve an importance score for each feature. The feature importance is calculated for a single decision tree by the amount that each features’s split point improves the performance, weighted by the number of observations the node is responsible for. Adopting this idea to our experimentation we use XGBoost’s feature importance to identify the most important DREMI scores for each of the developed cell type-specific models of survival. To account for reproducibility, we repeat the experimentation 1000 times, and we report the most frequently selected/stable features (31). Our results show (Figure 4b-h) the feature importance in association with the selection frequency.

We find the following top-1 cell type -specific features with the potential to discriminate STS from LTS patients within our cohort: pP38→pSTAT3 for B cells, pSTAT3→pCREB for CD4+ T cells, pCREB→pSTAT3 for CD8+ T cells, pSTAT3→pErk for Hematopoietic Stem Cells and Multipotent Progenitor cells (HSCs_MPPs), pErk→pSTAT5 for Monocytes, pS6→pSTAT5 for Natural Killer cells (NK) and p4EBP1→Casp3 for plasmocytoid Dendritic Cells (Supplementary Figure 13).

Importantly, our experimentation with DREMI scores and ML-based feature importance identify signalling dynamics (i.e., DREMI score between proteins) that are easier to interpret in a precision oncology application. To showcase their benefit for patient stratification, we perform independent univariate Kaplan-Meier (KP) survival analysis (Supplementary Figure 14a-g). All the selected DREMI scores expect for pSTAT3→pCREB in CD4_T cells and pS6→pSTAT5 in NK cells lead to statistically significant separation of patients. Within our cohort multivariate Cox proportional hazard modelling demonstrates that higher pP38→pSTAT3 DREMI scores in B cells and higher pErk→pSTAT5 DREMI scores in Monocytes are associated with shorter survival of leukemia patients (Supplementary Figure 14h).

Taken together, the proposed computational framework enables data-driven stratification of patients, by targeting single cell interactions between phosphoproteins. And this provides an enhanced stimulus for the development of more effective patient stratification methods based on single cell omics. Importantly, our results can be exploited in the future, to investigate cellular heterogeneity of cell populations. Our framework adds a new dimension to the analysis of CyTOF data, with the possibility to link signalling protein dynamics with genomic aberrations, and this idea will be explored in the next subsection.

#### 3.2.4 DREMI scores synergise with genetic information for improved patient stratification

Current patient stratification methods in leukemia utilise clinical data and genetic information at the time of diagnosis to guide clinical decisions. Thus, it is of great importance to optimise existing risk stratification approaches. To explore this possibility in our case study, we utilise combination of important DREMI features from the previous subsection with genetic information such as the FLT3-ITD mutational status.

For comparison purposes, we first stratify the cohort based on the FLT3-ITD mutational status alone and univariate KP survival analysis is performed. From Figure 5a, becomes apparent that there is no difference in survival between wild-type and mutated patients. Next, we combine the FLT3-ITD status with the most important cell type specific DREMI scores for B cells, Monocytes and HSCs & MPPs. In all cases (Figure 5b-d) the worst survival is achieved by the wild-type patients with the higher levels of the selected DREMI scores. Interestingly, with the addition of DREMI scores we can detect a subgroup of wild-type patients with improved survival that was not seen before.

**Figure 5:**
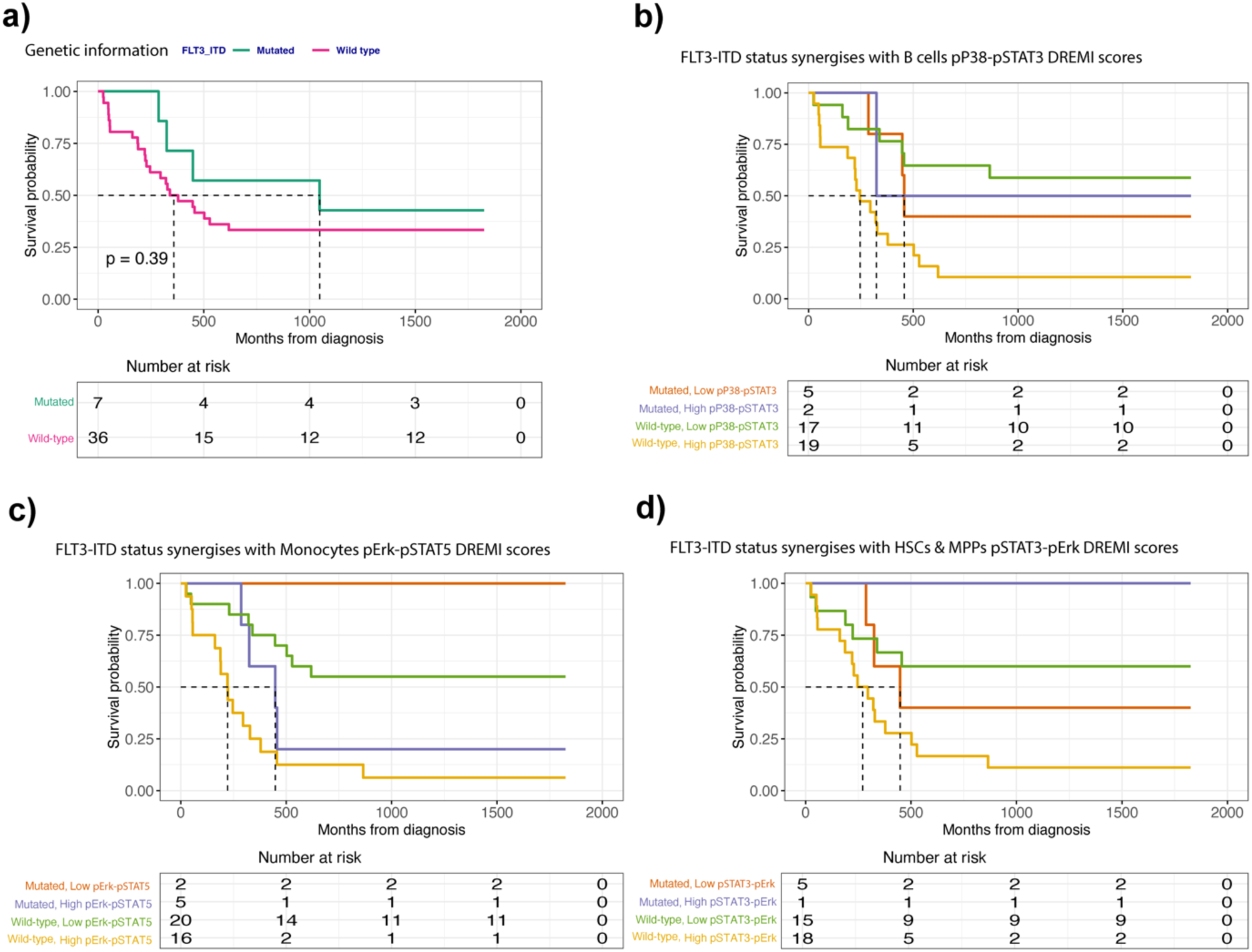
DREMI scores synergise with FLT3-ITD mutational status for risk stratification. a) The survival probability of all patients in the cohort with FLT3-ITD mutational status available (p=0.39); b) The survival probability of FLT3-ITD wild type (wt) and FLT3-ITD mutated patients combined with pP38-pSTAT3 DREMI scores computed from B cells;. c) The survival probability of FLT3-ITD wild type and FLT3-ITD mutated patients combined with pErk-pSTAT5 DREMI scores computed from Monocytes; d) The survival probability of FLT3-ITD wild type and FLT3-ITD mutated patients combined with pSTAT3-pErk DREMI scores computed from HSCs and MPP cells; Survival analysis is performed using Kaplan-Meier method.

Although our results are preliminary and limited by the small cohort size (i.e., seven mutated patients), we showcase that single cell signalling omics could complement genetic criteria such as hotspot mutational status, paving the way for improved risk stratification at time of diagnosis.

## Discussion

In this study we deployed a bioinformatics framework which facilitates automated cell type annotation guided by a reference dataset, enabling systematic exploration of protein dynamics for biomarker discoveries using CyTOF experiments.

Focusing on the cell type annotation task, the proposed approach is guided by a comprehensive reference dataset that is used to train different supervised and semi-supervised classification methods. Our self-consistency test and independent validations using several benchmark datasets show that the Scaffold approach is easy to implement, fast, and robust allowing for effective annotation of cells that are unseen from the reference cells used for training. The improved generalisation opens possibilities for automated and scalable data-driven exploration of single cell datasets in future projects.

Furthermore, to showcase the utilisation of the developed framework in a precision oncology application we perform a case study focusing a cohort of 43 leukemia patients. The Scaffold approach is used to ‘align’ cells from leukemia patients to the reference and facilitate cell type annotation. On top of the developed cell type annotation framework, we aim at predicting survival using ML-based approaches. To mitigate the limitation of single-value statistics (e.g., median protein expression values) for feature extraction, we introduce a novel feature engineering technique based on DREMI scores. The proposed feature engineering technique is based on mutual information, and it is agnostic of signalling network topologies that are frequently altered in disease conditions. Instead, it can be applied to quantify the strength of information transfer between all possible pairs of (e.g., edges in a signalling network topology) the available proteins in a CyTOF antibody panel. DREMI-derived features are in principle easier to interpret in a biological context since they consider how information is transmitted between cells in a signaling cascade. Thus, unique opportunities are opened to discover novel signalling dynamics that are predictive of the disease outcome.

Following this rational we compare different ML classification algorithms and find that the XGBoost algorithm surpasses the prevalent regularised regression methods namely LASSO and Ridge for survival prediction. Most importantly, we assess the impact of the class imbalance problem in survival prediction which was neglected by similar studies. Our results re-confirmed the effectiveness of SMOTE for imbalanced machine learning. We also utilise the XGBoost’s feature selection component to identify salient signalling dynamics that are predictive of survival. Furthermore, the identified cell type-specific features are used in combination with ‘standard’ genetic information such as the FLT3-ITD mutational status allowing for improved patient stratification. Our results highlight the translational potential of the developed method that can be readily used to combine signalling states measured with CyTOF with ‘standard’ clinical and genetic information for patient stratification. Importantly, the bioinformatics tools and the analysis paradigm presented here, can be useful in the future to investigate mechanisms of therapy resistance and possibly identify vulnerabilities that could serve as novel therapeutic targets.

Focusing on possible limitations of this work, the effectiveness of the Scaffold approach depends on the quality of reference dataset. It also depends on the antibody panel compatibility between the reference dataset and the dataset to be annotated/tested. Within our experimentation we were only able to detect seven cell types, and we were limited on predicting less prevalent and ‘rare’ cell populations that require more specific cell type defining antibodies. Thus, it is recommended to study carefully the compatibility of the datasets before annotation and focus on the most appropriate subsets of the available cell types in the reference.

There is also room for improvements targeting the ML modelling part presented in our case study. Similar to other studies, the limited cohort size restricts the generalisation of ML-based findings. Thus, more work is required to assess the validity of findings in larger cohorts of patients. For example, it will be interesting to test the proposed methodology to larger cohorts of leukemia patients. However, it is not straightforward to generate or find and download diagnostic CyTOF samples with compatible antibody panels, that match the demographic and clinical characteristics of our leukemia cohort. In addition, with different antibody panels (e.g., surface phenotypic markers and/or signalling phosphoproteins), we anticipate that the proposed computational framework could easily be extended to investigate different cell types and signalling proteins (32). There also improvements targeting the implementation of a graphical-user interface (GUI) with visualisation functions for the Scaffold approach to make cell type annotation interactive. In this way, it would be easier for users with no prior programming knowledge skills to visualise immune landscapes and explore the structure of their datasets facilitating comparisons with the reference. Finally, utilising existing knowledge from annotated datasets following the developed framework is a promising research direction that can be further exploited in the future with more sophisticated reference-guided cell type annotation algorithms (19).

Focusing on data interpretation and translational medicine aspects, the development of predictive models of survival could benefit from integration of additional single cell omics. Our results demonstrate that it is possible to incorporate single cell omics in combination with ‘standard’ clinical and genetic characteristics. For example, in future studies state-of-the-art classification systems for leukemia such as the European LeukemiaNet (ELN) risk classification system for Acute Myeloid Leukemia (33), or the PAM50 classification algorithm for breast cancer could be complemented with information from CyTOF, imaging mass cytometry (IMC) or spatial transcriptomic datasets that capture the cellular heterogeneity of patients as well as aberrant signalling states that are disease-specific (34). It is also worth noting that the developed framework although it has been originally designed to predict survival in leukemia using CyTOF, it can be in principle applied to different diseases and single cell technologies as well.

Collectively, our study streamlined the application of different ML algorithms on CyTOF data to annotate cell populations and engineer features for patient stratification with ML. We believe that the utilisation and exploration of single cell data, as described here, could provide a more general bioinformatics framework with many applications in future precision medicine projects.

## Methods

### Publicly available datasets used in the study

For the development and evaluation of the cell type annotation part of our framework we use the following publicly available datasets:

1. **Reference dataset:** we downloaded a multimodal reference dataset acquired from Abseq experiments measuring simultaneously 97 surface proteins (e.g., antibodies) and 462 mRNAs in 49,057 cells from healthy blood and bone marrow (26). In its original publication the dataset was annotated using multi-omics computational approaches and 43 different cell types were discovered. In our study, we exclude cell types with less than 300 cells which results to 31 cell types after filtering. In Supplementary Figure 1, we use the original UMAP coordinates to visualise the 2D structure of the data, accompanied with other summary statistics about the relative abundance of the populations found in this dataset. During our experimentation this dataset is used for training different computational approaches (see sections below) as well as for ‘guiding’ the annotation of unknown cells using the Scaffold approach.
2. **Benchmark datasets:** we downloaded three different datasets that are used as ‘ground-truth’ to assess the performance of different ‘automated’ methods for cell type annotation (6,11). We use: a) the AML dataset (referred here as AML_benchmark) that is a healthy human bone marrow mass cytometry dataset consisting of 104,184 cells screened with 32 markers resulting in 14 cell populations defined by manual gating; b) the BMMC dataset (refereed here as BMMC_benchmark) which is also a healthy human bone marrow dataset, consisting of 81,747 cells screened with 13 markers, resulting to 24 cell populations defined by manual gating; c) finally, we use the ‘Samusik_01’ dataset containing 53,173 cells (referred here as PANORAMA_benchmark although it is fraction of the full PANORAMA dataset), of mice bone marrow, screened with a panel of 39 markers and manually gated into 24 cell populations. During our experimentation and to harmonise differences between the antibody panels of the reference and the benchmark datasets, we focus on small set of seven cell populations namely Monocytes, NK, B-cells, CD4_T, CD8_T, pDC, HSC_MPP (Supplementary Figures 3-8). During our experimentation the benchmark datasets are used for testing, but the training is performed using the reference dataset from Triana et al.. Although this scenario is extreme for ML modelling, it reflects the pragmatic scenario of reusing existing knowledge for annotating cell populations.

To showcase the use of the developed framework in a precision medicine application, we downloaded data from a leukemia cohort (21). To increase cohort size, our case study is top-ed up with additional leukemia samples that were collected and processed in the same batch together with the published data from ref. 21, resulting in total to 43 peripheral blood samples collected at time of diagnosis. More details about the demographics of the published cohort can be found in its’ original publication (patient numbers from the original publication are shown). Supplementary Table 1 and Supplementary Table 2 summarise patient characteristics for the extended leukemia cohort we used. Mass cytometry experiments for all samples were originally performed using an antibody panel of 21 surface proteins and 15 signalling proteins that was designed and optimised in ref. 21. In total 6,425,200 single cells were screened and used in our case study. For comparison purposes we also downloaded a cohort of seven healthy donors resulting to 1,242,733 single cells that were screened with the same antibody panel as part of the same leukemia study (21). During our experimentation, the leukemia cohort is annotated using the Scaffold approach. The Granulocytic population (cells low in CD45 expression and high for CD66b expression) is excluded from our analysis since it was not matching the reference dataset (CD66b marker is absent from the backbone). Then, cell type-specific data are subjected to ML modelling to identify predictors of survival.

### Computational approaches for cell type annotation

To benchmark the performance of different methods for cell type annotation we utilise two state-of-the-art supervised classification algorithms namely the K-Nearest Neighbours (KNN) and the Linear Discriminant Analysis (LDA), a recently published semi-supervised approach called CyAnno (15) as well as an adaptation of the Scaffold approach introduced by ref. 27.

Given an unseen cell named c1, a supervised classifier for cell type annotation makes a prediction by assigning c1 to cell type Ci (e.g., in our case study i ε [Monocytes, NK, B-cells, CD4_T, CD8_T, pDC, HSC_MPP]). LDA considers the posterior probability of c1 to be maximized, across the available cell types. KNN on the other hand, computes the distance of c1 to k nearest neighbors, and applies a majority voting technique to assign cell types. The original publication by Abdelaal et al. applied a posterior probability threshold to reduce misclassification error of LDA (11). However, in our study we did not apply this technique. For the KNN classifier to assess the impact of parameter k, we repeated use of KNN with values K=3 and K=10.

CyAnno on the other hand is a semi-supervised approach that requires a manually gated training set to identify ‘landmarks’ (defined as smaller subsets of representative cells), that are used to build ‘one-vs-rest’ cell type specific binary classification models. In our experimentation since we did not have manually gated data, we execute CyAnno providing as input annotated cell types from the reference dataset.

Finally, we adopt the concept of the Scaffold approach to map unseen cells to the reference dataset. The Scaffold approach is implemented in-house adopting codes from the grappolo and vite R packages. Scaffold mapping for unseen data works as follows: a) Single cells are first clustered using the FlowSOM algorithm (9) using a grid of size 10x10 as a default option; b) For each cluster, the centroids, which are scaffold’s landmarks, are computed using selected ‘backbone’ surface markers. In our experimentation ‘backbone’ markers for each dataset are the common markers between the testing datasets and the reference; c) The landmarks from the unseen dataset are then compared with all landmarks from the annotated reference dataset and their cosine similarity is computed. The original Scaffold implementation uses force-directed graphs to guide data organization into the 2D space. Nodes that are clusters of cells in our case, are connected by edges with a length proportional to their cosine similarity. These nodes can be spatialized into a graph and visualised to understand the structure and the relationships of cell types in the immune system; d) in our case since the aim is only to automatically assign cell populations, we do not visualise the immune system as graph, but we retrieve the edges with the highest weights. The reference landmark node of the highest weight edge is subsequently used to assign the cell types.

### Generating features for ML-based modelling with CyTOF

#### 1. Features generated using median expression values of proteins

To generate features for ML-based modelling with CyTOF, we first use the median intensity values of phosphoproteins in the antibody panel. In our case study we generate feature vectors of 15 variables describing the expression levels of the following markers: pAxl, CyclinB1, pNFkB, pErk, pSTAT1, pP38, pSTAT3, pCREB, pHist3, Casp3, pSTAT5, p4EBP1, pAkt, pRB, pS6. Median expression values as features for ML-based modelling provide a fast and easy to implement solution, however, do not take full advantage of the rich single cell expression distributions of proteins measured with CyTOF. Technical noise, natural stochasticity, and many other factors have collectively confounding effects on the analysis of signalling proteins using single-value statistics. Although we are fully aware of the above-mentioned limitation, we decided to use median-derived features as a baseline approach for our experimentation.

#### 2. Features engineered using DREMI scores

To mitigate the limitations of single-value statistics for engineering features from CyTOF, we utilize DREMI that was originally introduced in ref. 25. A key conceptual advantage of DREMI compared to other approaches, is that DREMI computes mutual information based on resampled conditional density estimations. Given proteins X and Y in a network/pathway with the edge X→Y, DREMI quantifies how the state of Y varies with different states of X, which is different from describing the density of cells states X and Y in a conventional scatter plot or in a joint density plot. In other words, DREMI reveals how protein Y changes as a function of protein’s X activity, providing a quantitate score to characterise the ‘strength’ of this signalling relationship.

Generating feature vectors using DREMI, one could target few proteins that are known to interact in a signalling cascade, or in an agnostic manner by evaluating all possible pairwise combinations of proteins in the antibody panel. The latter approach is applied in our study, considering both signalling protein directions (e.g. X → Y and Y → X), and across the studied cell types. We use a minimum number of n=200 cells for DREMI estimation per cell type, and thus we exclude from the analysis patients with low cellular abundance for some cell types. DREMI was implemented originally in MATLAB, but the codes are re-written in R and incorporated within our developed framework. In summary, 210 features using all possible combinations of 15 available signalling proteins in the panel are generated considering both directions of signalling relationships.

### Predicting patients’ survival using machine learning

The problem of predicting patients’ survival is formulated as a binary classification task to discriminate between patients of short-term survival (STS) versus long-term survival (LTS). In our case study, a period of 60 months after diagnosis was used by clinicians to evaluate survival’s status. Based on the available samples and clinical information, n=15 patients are in LTS group and n=28 patients are in STS group. To predict survival our framework deploys cell type-specific ML modelling using automated cell type annotations from the previous part (Supplementary Figure 11). We also test two different feature sets, one generated using medians (n=15 features) and another using DREMI score (n=210 features) as described in the previous subsection.

We test three different classification algorithms namely the XGBoost from the eXtreme Gradient Boosting R package, as well as LASSO regression and Ridge regression from the glmnet R package. To note that for both LASSO and Ridge implementations the binomial family function is used with parameter alpha=0 for LASSO and alpha=1 for Ridge.

To account for the class imbalance problem between positive and negative classes in the dataset, we use the SMOTE implementation of the DMwR R package. With SMOTE the ML modelling is repeated using synthetic datasets with different number of instances from the majority and minority classes. We choose SMOTE’s internal parameters perc.over and perc.under so as ratios 1:1, 1:2 and 1:3 between LTS and STS classes are generated.

Finally, to assess the classification performance and optimise the internal parameters of the classification algorithm the hold-out approach is deployed. To do so, the available data instances are split randomly into two disjoint sets one used for training and validation and the other for testing. To account for reproducibility the random split of the samples is repeated 100 times and every time an independent cell type-specific ML classifier is trained/validated and tested (e.g., in total 700 classifiers are developed for the original data (100*7 cell types) and 3*700 classifiers are developed for SMOTE data). We also use two different random splits of the data: one split with 80% of the available instances for training/validation, and the remaining 20% for testing; and another split with 50% of the available instances for training/validation and the remaining 50% for testing. Throughout our experimentation the validation subset is the 40% of the available instances of the training/validation set.

All algorithms are tuned using the validation dataset during the algorithms’ training process. For the XGBoost algorithm fine tuning is performed for the maximum depth of a tree in the range of values [2,4,6,8,10,12,14,16,18,20,30,40,50,60]. For LASSO and Ridge, we consider the lambda value. In all cases the objective is to maximise F1 score in the validation set. A snapshot of parameters’ fine tuning is shown in Supplementary Figure 15.

### Performance metrics used in the study

We use different metrics to assess the performance of the developed ML models. In our study, the problem of classifying cells to different cell types is a multi-class classification problem, and thus the performance is assessed for every cell type separately using the ‘one-vs-all’ approach. Using this conversion, we generate 2x2 confusion matrices per cell type and the performance is computed using the following metrics:

1. 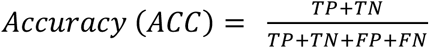
2. 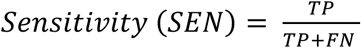
3. 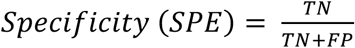
4. 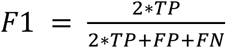

Where, TP denotes true positive, TN denotes true negative, FP denotes false positive, and FN denotes false negative predictions.

Next, the problem of predicting short-term survival from long-term survival is formulated as a binary classification task and the performance is assessed using the standard 2x2 confusion matrix and metrics Sensitivity, Specificty and F1 using the previous formulas. We also use the Area Under Curve (AUC) implemented using the roc R package.

### Statistical analysis

Statistical analyses and tests are implemented in the R package. To test statistically significant differences between abundance distributions of two groups, we use Wilcoxon Rank Sum test, or Anova with TukeyHSD correction for more than two groups when applicable.

Patients are stratified to low and high groups of DREMI scores using the median DREMI score estimated from the full set of cohort values.

Kaplan-Meier analysis is used to estimate survival rates. Differences between different groups of survival are assessed using log-rank test. The Cox proportional hazards model is used to investigate the association between the survival time of patients and one or more input variables such as cellular abundance or DREMI scores. The assumptions of Cox proportional hazards modeling are tested using statistical tests and graphical diagnostics based on the scaled Schoenfeld residuals.

### Code and data availability

The framework has been implemented using the R language. The codes along with how to use instructions are available in GitHub https://github.com/dkleftogi/singleCellClassification. All datasets used in the study are publicly available and links to the original data sources and re-analysed data are provided in our GitHub page and in our dedicated data archive https://zenodo.org/records/10984478.

## Acknowledgements

The authors thank nurses and physicians in Bergen and Oslo, the pateint’s organization Blodkreftforeningen for following and supporting the project, personnel at the Flow Cytometry Core Facility at the University of Bergen and personnel at the Gjertsen- and Enserink labs for their skillful technical assistance. A special thanks to the participating patients and their families who suffered from leukemia and nonetheless gave the gift of participation so that others might benefit.

## Author Contribution

B.S.T, M.H, O.F, I.K.F.M, S.E.G, A.L and Y.F designed CyTOF experiments, performed the experiments, and provided input regarding the clinical data used in the leukemia case study; D.K developed the algorithms, and analysed the data; N.V.D.M, E.G, J.J.S, B.T.G, I.J., and G.S. provided intellectual input and help edit/revise the manuscript; D.K conceived the idea and led the project; D.K wrote the manuscript.

## Consent for publication

Patients have consented that mass cytometry data with no personal identifiers can be used for publication in an anonymized format.

## Supplementary Figures

**Supplementary Figure 1:**
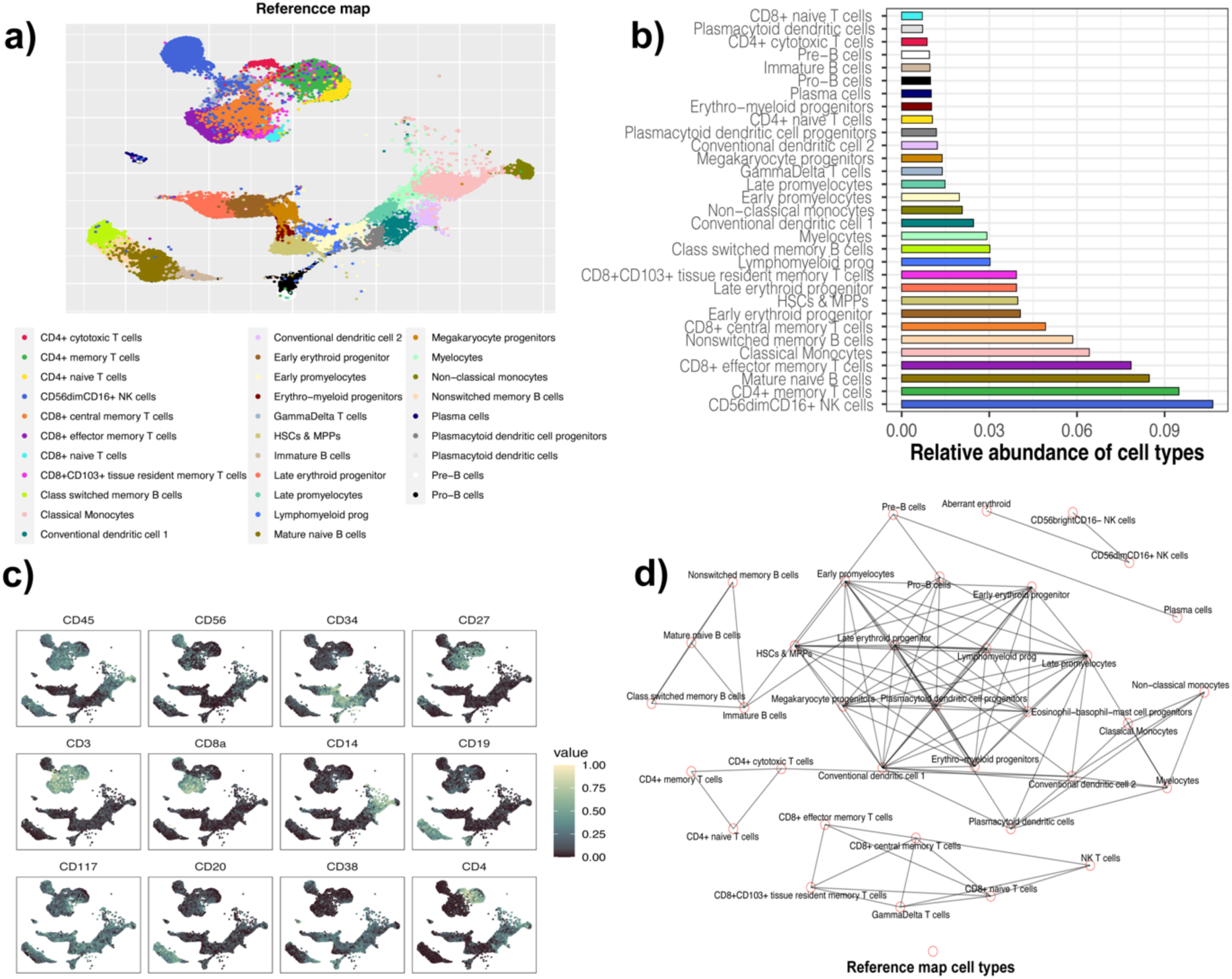
Overview of the annotated dataset used as reference. a) Original UMAP representation of the annotated reference dataset showing 31 cell types with more than 300 cells available; b) Barplot showing the relative abundances of 31 cell types found in the reference dataset; c) Original UMAP representation of the annotated reference dataset with overlaid expression of selected cell type defining markers; d) Force-directed graph generated using the Scaffold map approach. Nodes represent all cell types found in the reference dataset (43 in total without filtering of 300 cells), and edges indicate similarity between the cell types. A cosine similarity cut-off of 0.75 has been used to remove edges between nodes.

**Supplementary Figure 2:**
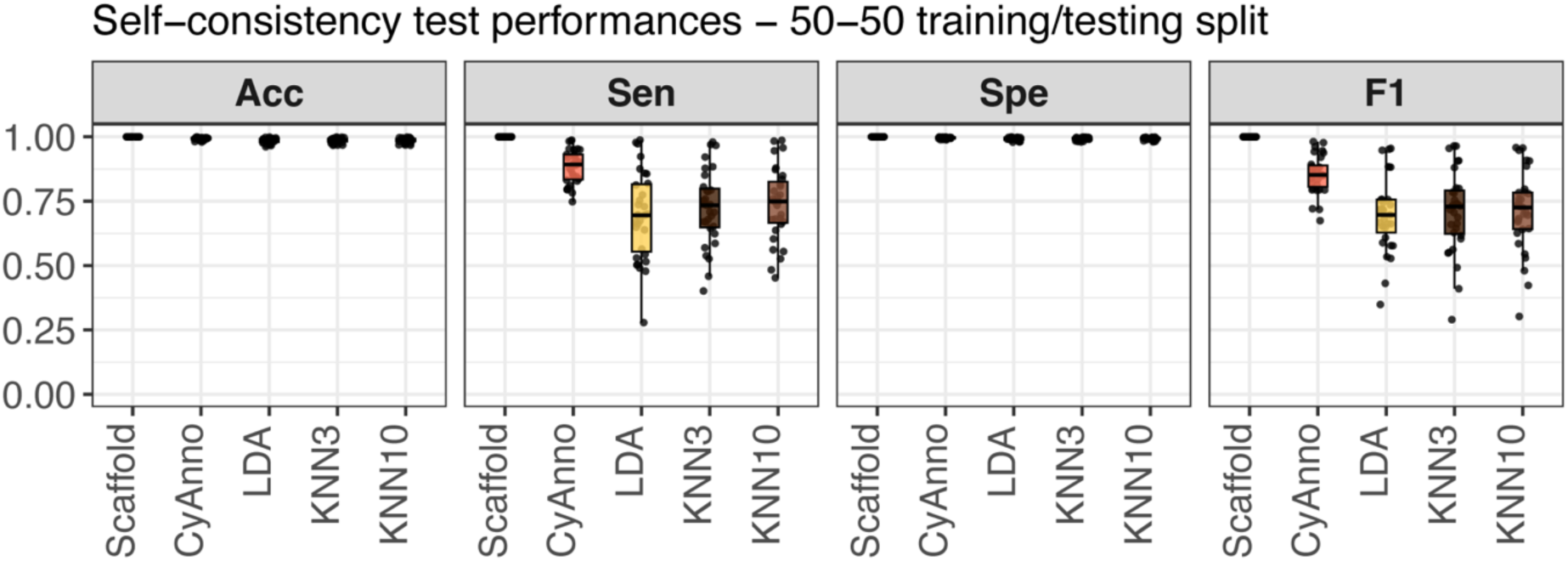
Performance evaluation using a self-consistency test. Results summarising the classification performance of Scaffold, CyAnno, LDA and KNN with K=3 and KNN with K=10, across 31 cell types from the Reference dataset, when split 50-50 is applied to generate disjoint sets for training and testing.

**Supplementary Figure 3:**
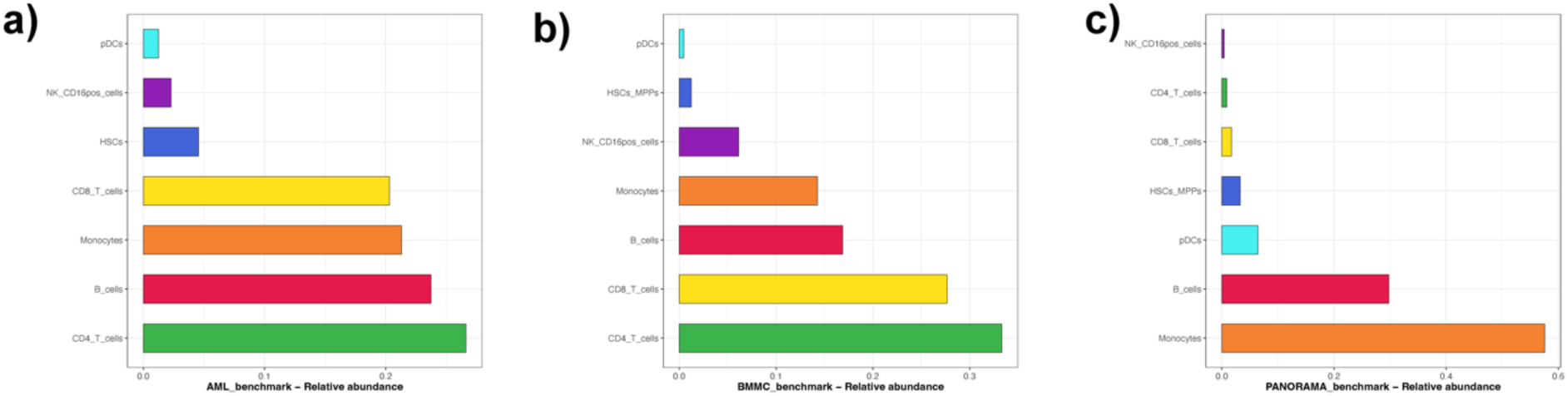
Overview of mass cytometry datasets used as benchmarks for cell type annotation. Barplot showing the relative abundance of cell types found in: a) AML_benchmark; b) BMMC_benchmark; and c) PANORAMA_benchmark.

**Supplementary Figure 4:**
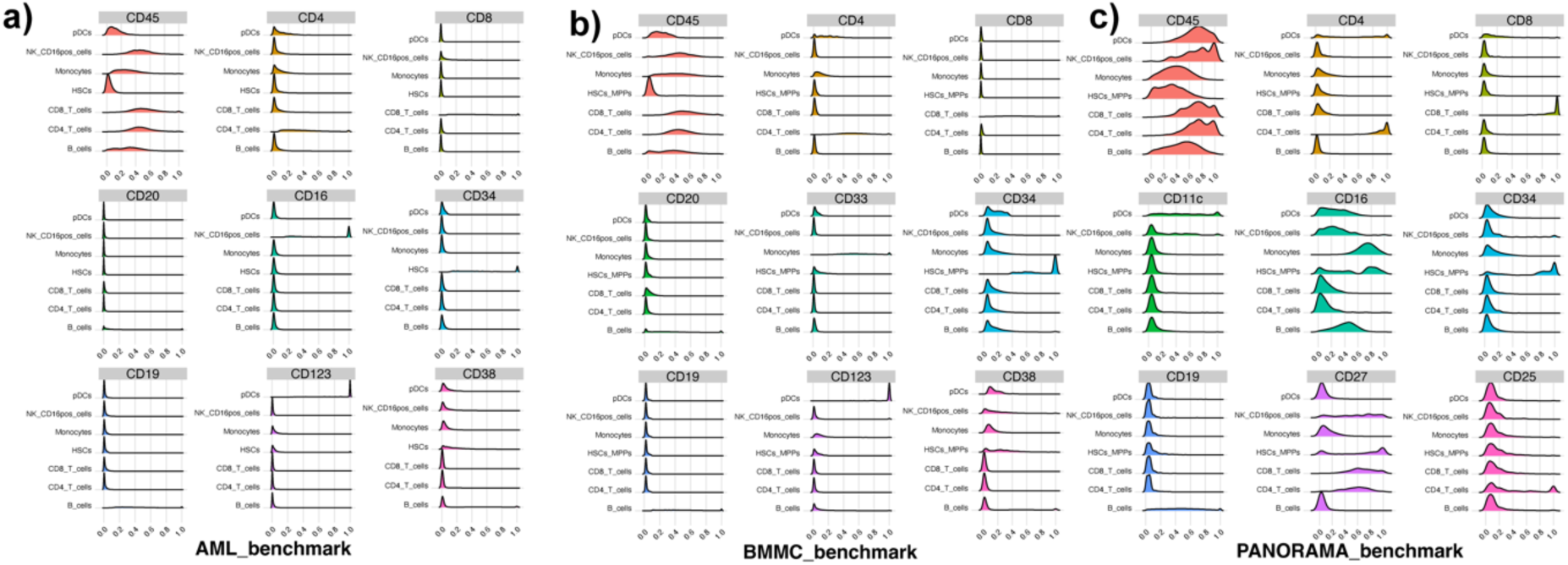
Inspecting marker expression profiles of the benchmark datasets. Ridge plots showing the expression profiles of selected cell type defining markers for seven cell types found in: a) AML_benchmark; b) BMMC_benchmark; and c) PANORAMA_benchmark. For visualization purposes, the expression values were scaled between 0 and 1 using low (1%) and high (99%) percentiles as boundaries. The x-axis shows the density of the scaled expression.

**Supplementary Figure 5:**
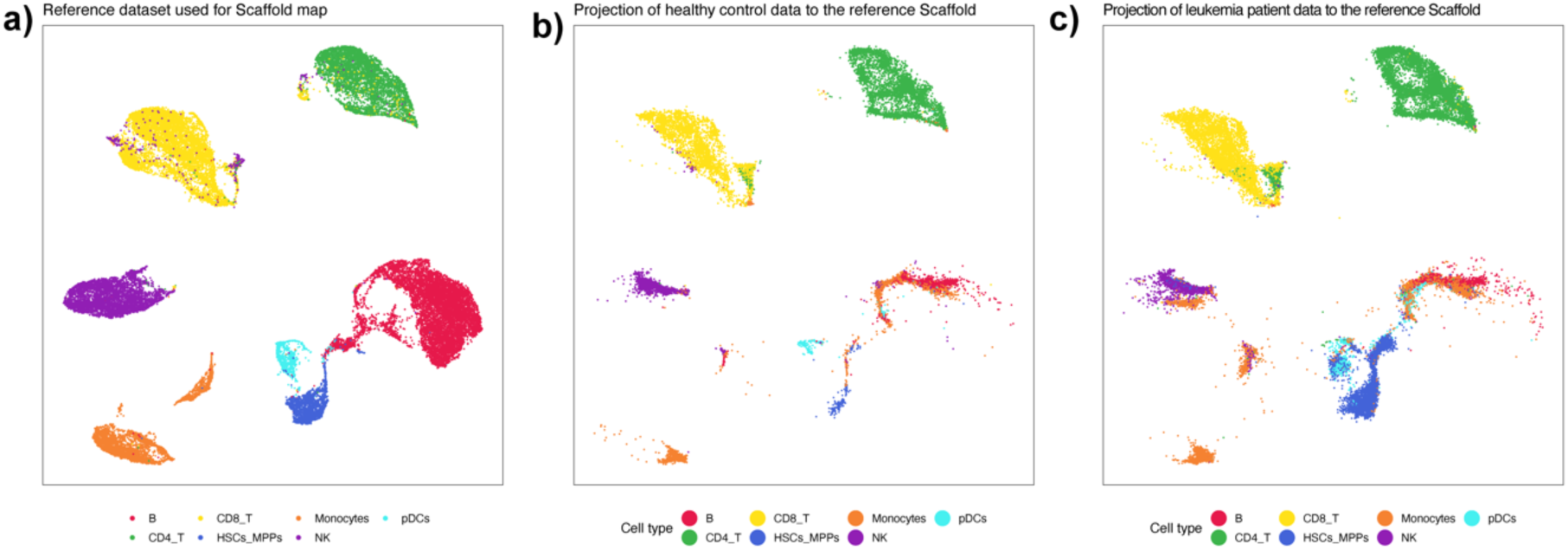
UMAP projections to the reference dataset. a) UMAP visualisation of the reference dataset constructed using a smaller set of ‘backbone’ markers namely: CD11b, CD8a, CD33, CD34, CD3, CD123, CD56, CD14, CD117, CD38, CD4, CD16, CD20, CD45, CD7 ; b) UMAP projection of the healthy control dataset to the reference dataset constructed using the same set of ‘backbone’ markers; c) UMAP projection of the leukemia cohort to the reference dataset constructed using the same set of ‘backbone’ markers.

**Supplementary Figure 6:**
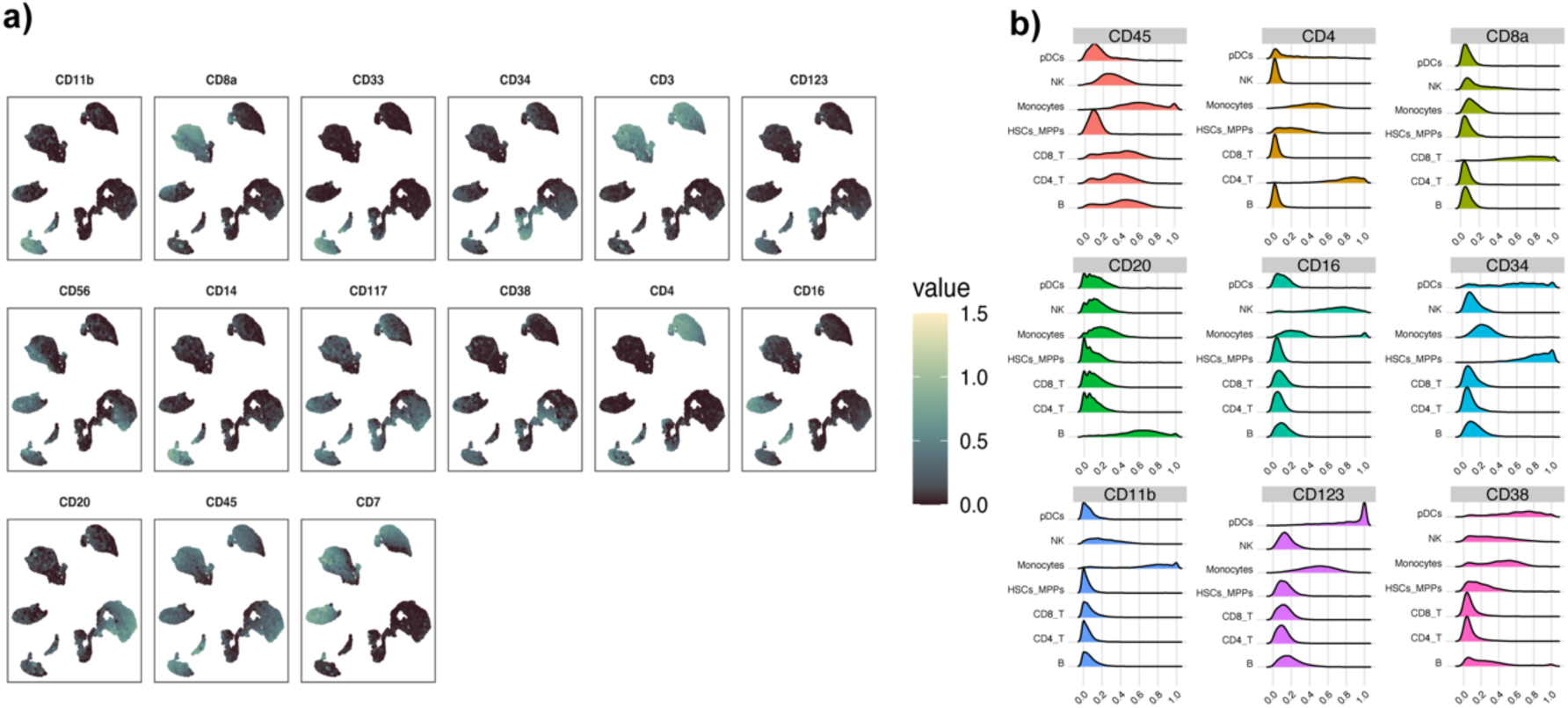
Inspecting marker expression profiles of the annotated AML_benchmark. a) UMAP representation of the annotated dataset with overlaid expression of selected cell type defining markers; b) Ridge plots showing the expression profiles of selected cell type defining markers for the predicted cell types.

**Supplementary Figure 7:**
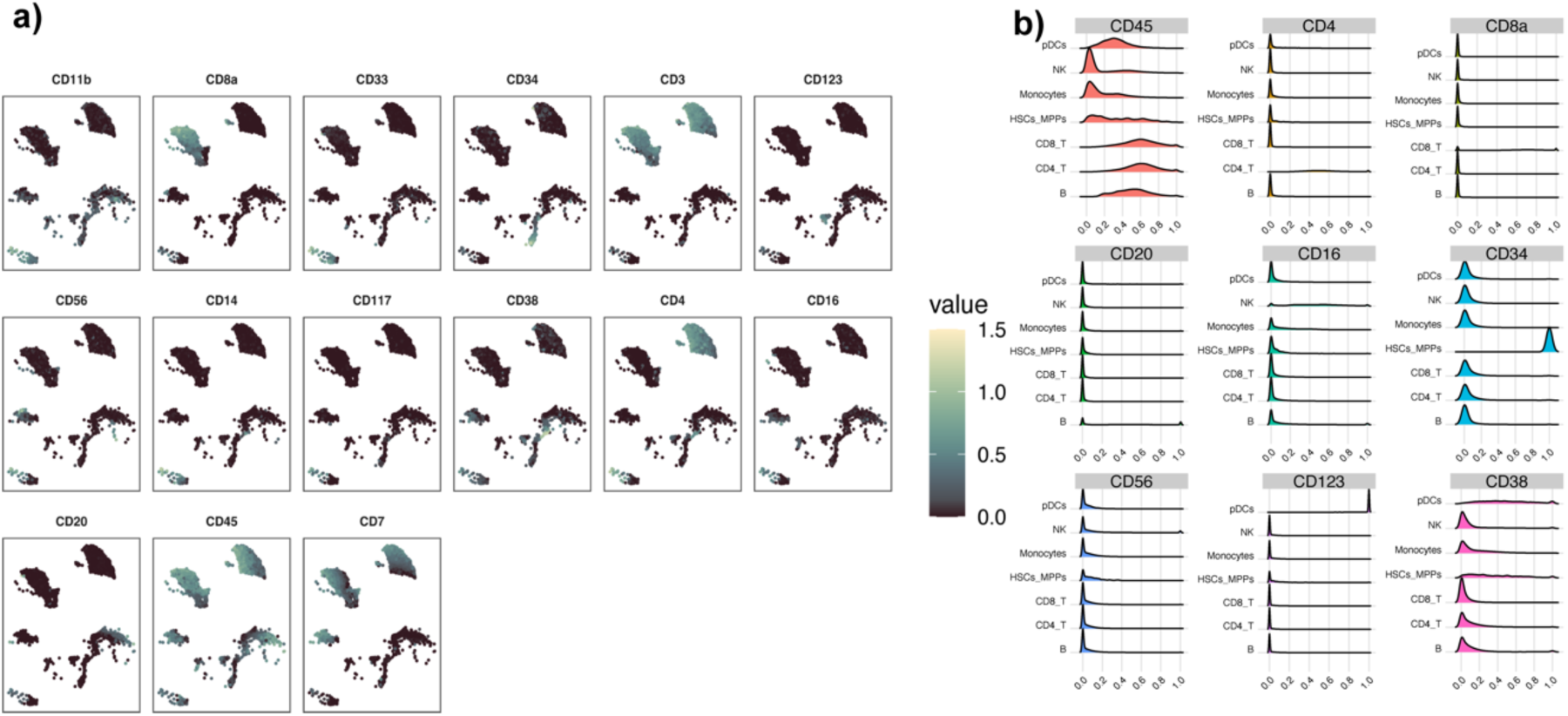
Inspecting marker expression profiles of the annotated BMMC_benchmark. a) UMAP representation of the annotated dataset with overlaid expression of selected cell type defining markers; b) Ridge plots showing the expression profiles of selected cell type defining markers for the predicted cell types.

**Supplementary Figure 8:**
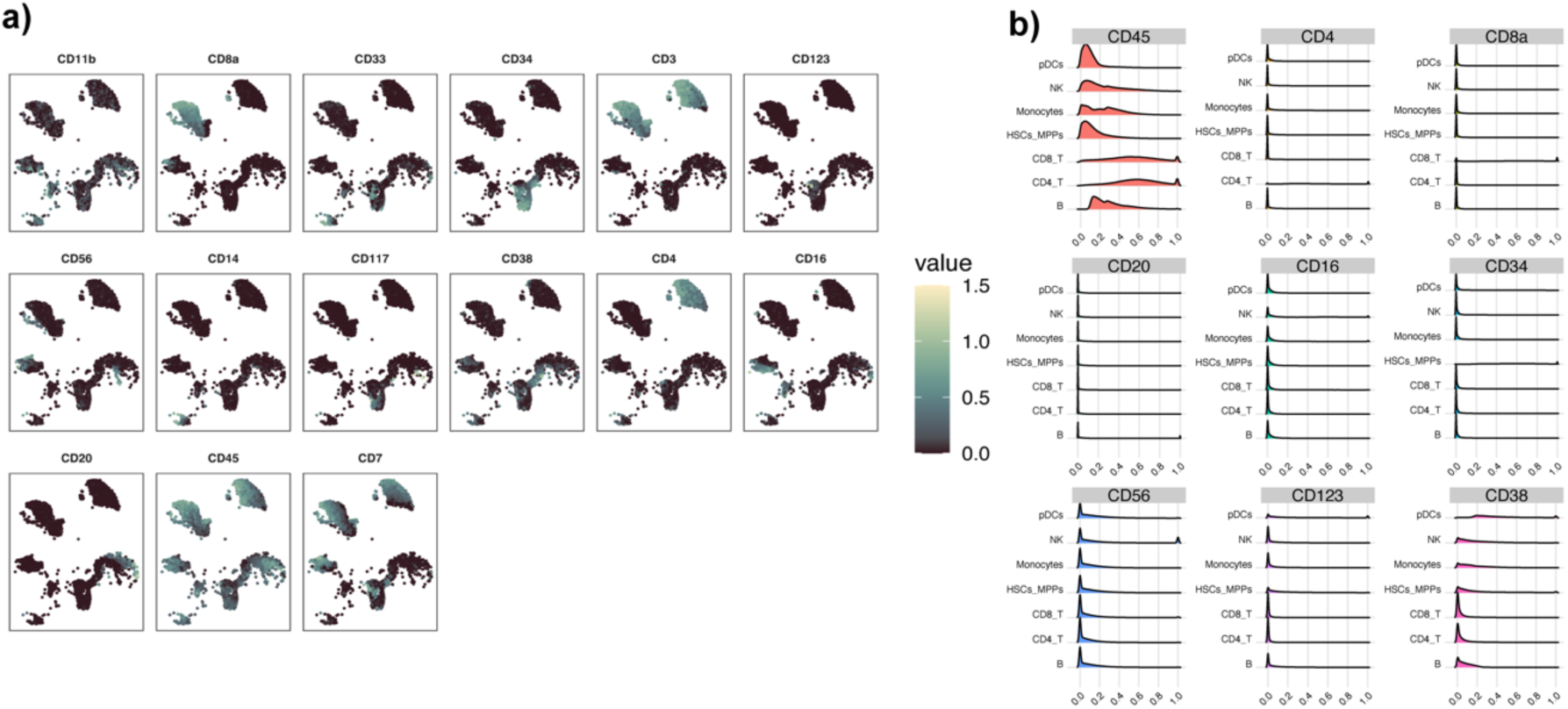
Inspecting marker expression profiles of the annotated PANORAMA_benchmark. a) UMAP representation of the annotated dataset with overlaid expression of selected cell type defining markers; b) Ridge plots showing the expression profiles of selected cell type defining markers for the predicted cell types found in PANORAMA_benchmark.

**Supplementary Figure 9:**
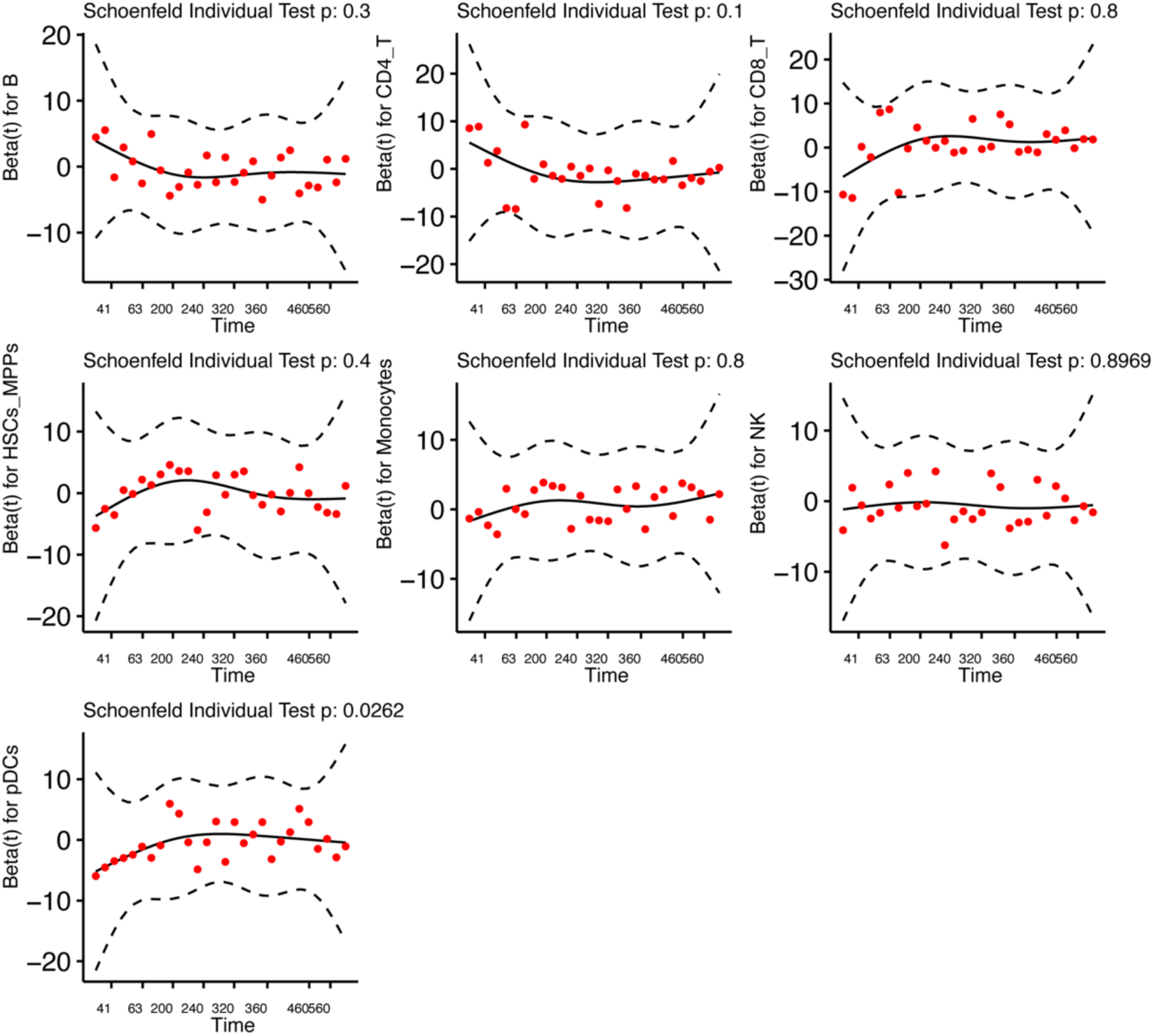
Visualising Schoenfeld residuals for multivariate Cox proportional hazard modelling.

**Supplementary Figure 10:**
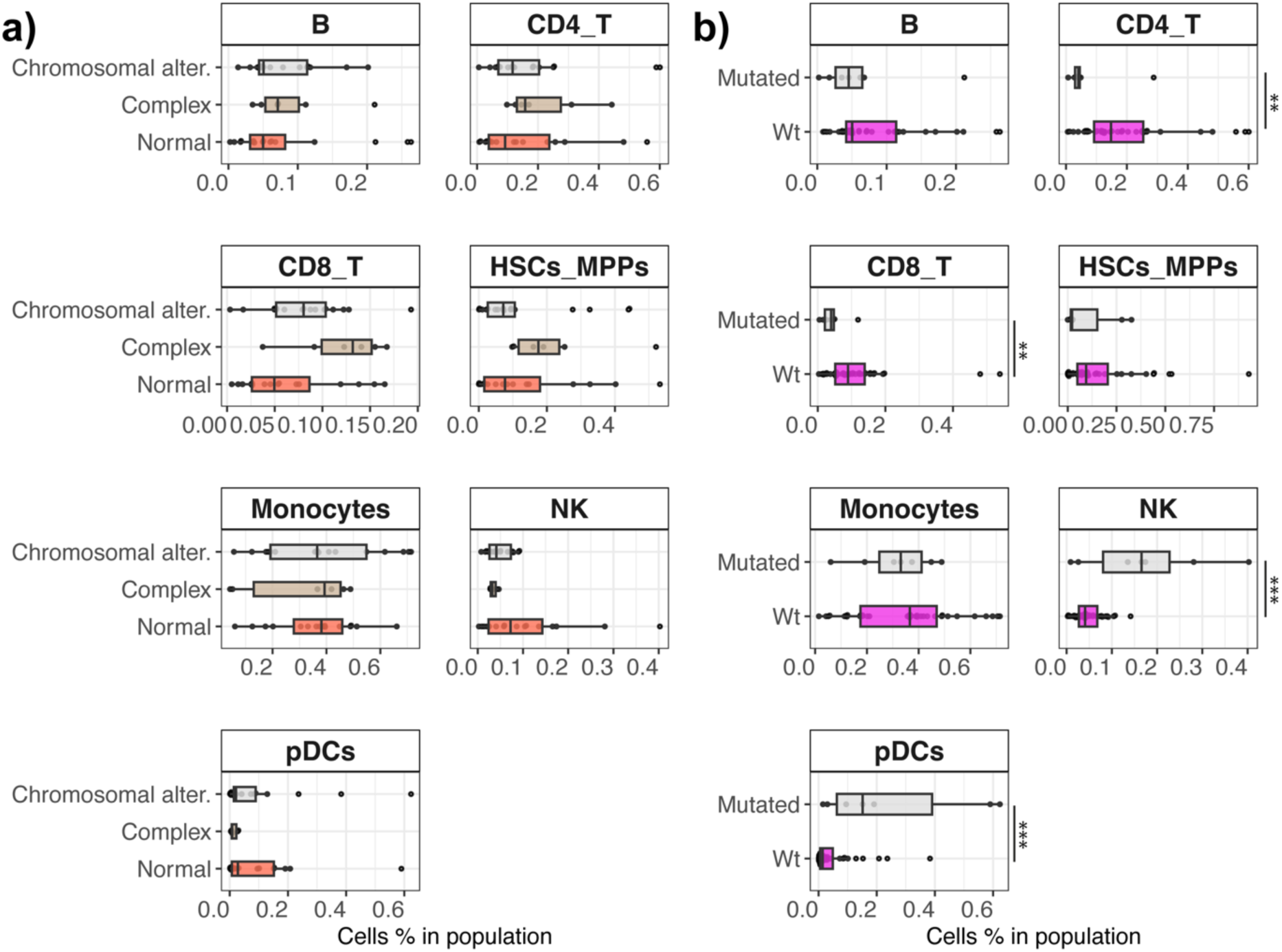
Comparing cellular abundance distributions of leukemia patients stratified by genetic information. a) Boxplots showing the cellular abundance of patients with karyotypes classified as normal, complex, and chromosomally altered; b) Boxplots showing the cellular abundance of patients with FLT3-ITD mutated and FLT3-ITD wild type status. Statistically significant differences in a) were tested using anova with TukeyHSD correction, and in b) using Wilcoxon rank sum test. The levels of significance are: * for p<0.05, ** for p<0.01 and *** for p<0.001.

**Supplementary Figure 11:**
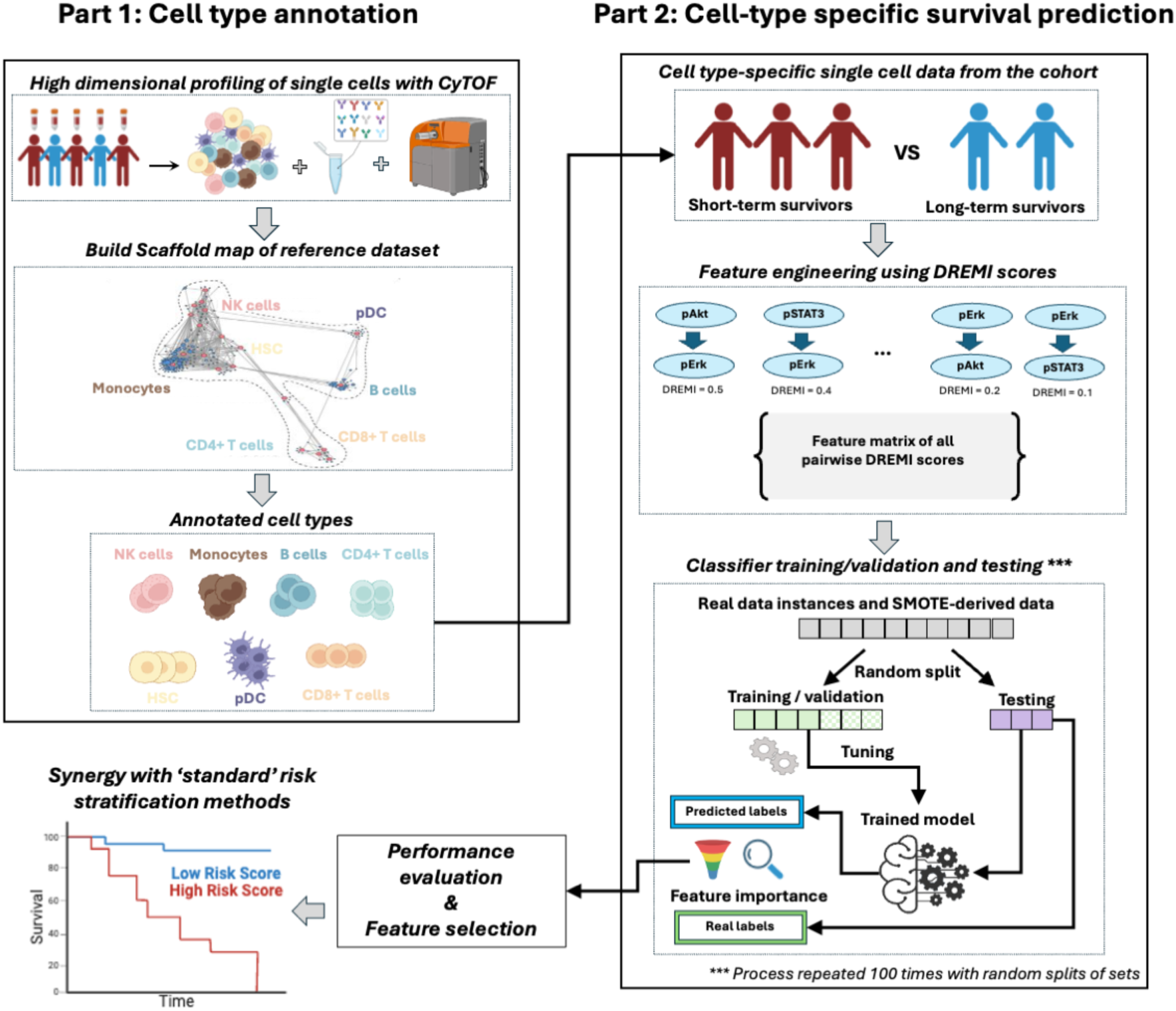
The workflow of the developed bioinformatics framework. Part 1 illustrates the cell type annotation method using the Scaffold map approach, and Part 2 illustrates the ML-based prediction of survival.

**Supplementary Figure 12:**
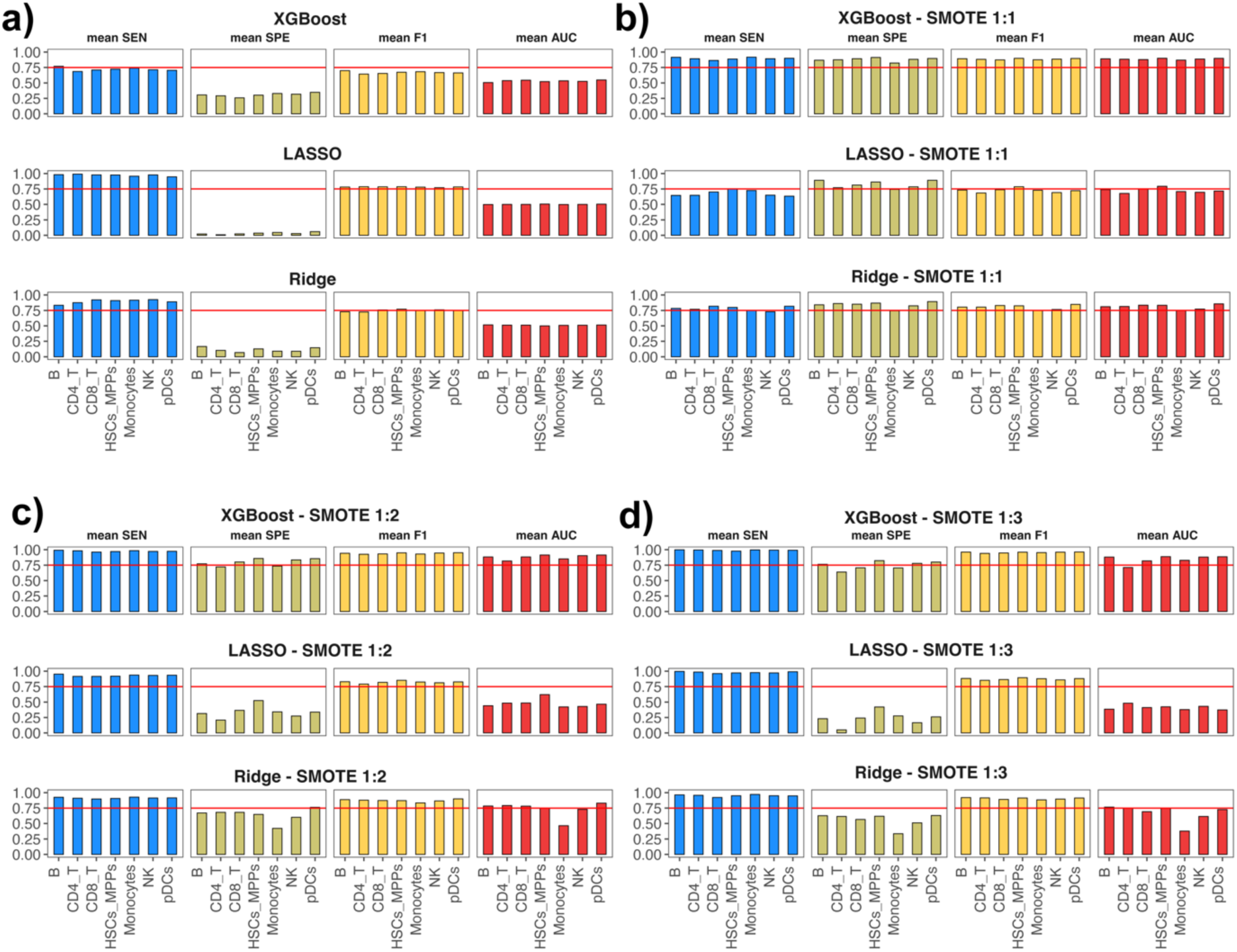
Performance evaluation of different ML-based methods for predicting survival using as input features median expression values of phosphoproteins. For all experiments 50-50 random split of training/validation and testing sets is performed and average classification performance of XGBoost, LASSO and Ridge regression for discriminating STS from LTS patients is shown for: a) the data from the original case study cohort; b) SMOTE-derived synthetic data with ratio 1:1 between the classes; b) SMOTE-derived synthetic data with ratio 1:2 between the classes; c) SMOTE-derived synthetic data with ratio 1:3 between the classes. The performance is assessed using mean Sensitivity (SEN), mean Specificity (SPE), mean F1 and mean Area Under Curve (AUC) of 100 executions.

**Supplementary Figure 13:**
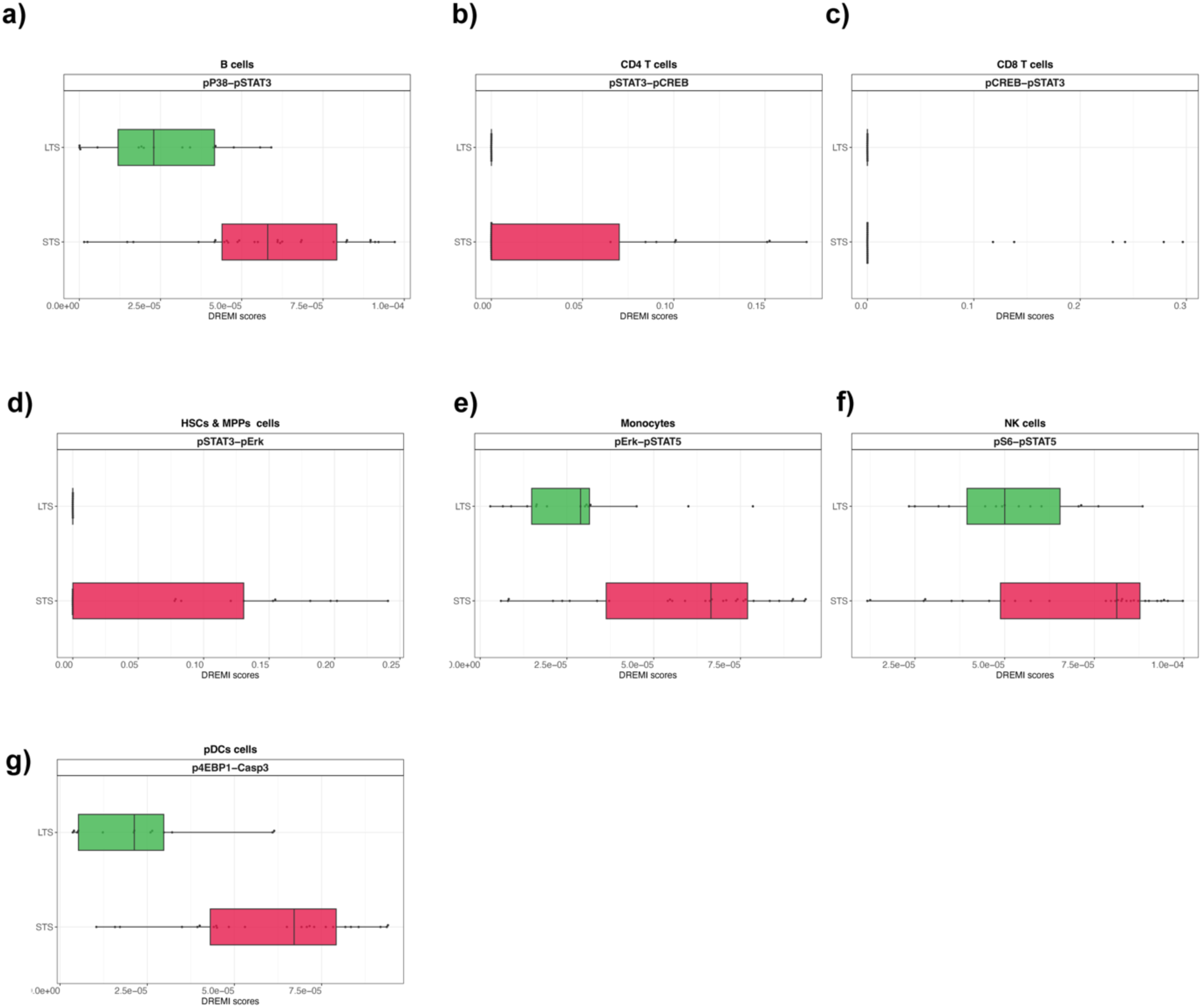
Visualising the distribution of the most important DREMI scores learnt from the XGBoost models. a) Boxplots showing B cell derived DREMI scores of pP38-pSTAT3 for STS and LTS patients; b) Boxplots showing CD4_T cell derived DREMI scores of pSTAT3-pCREB for STS and LTS patients; c) Boxplots showing CD8_T cell derived DREMI scores of pCREB-pSTAT3 for STS and LTS patients; d) Boxplots showing HSC_MPP cell derived DREMI scores of pSTAT3-pErk for STS and LTS patients; e) Boxplots showing Monocyte derived DREMI scores of pErk-pSTAT5 for STS and LTS patients; f) Boxplots showing NK cell derived DREMI scores of pS6-pSTAT5 for STS and LTS patients; g) Boxplots showing pDCs derived DREMI scores of p4EBP1-Casp3 for STS and LTS patients.

**Supplementary Figure 14:**
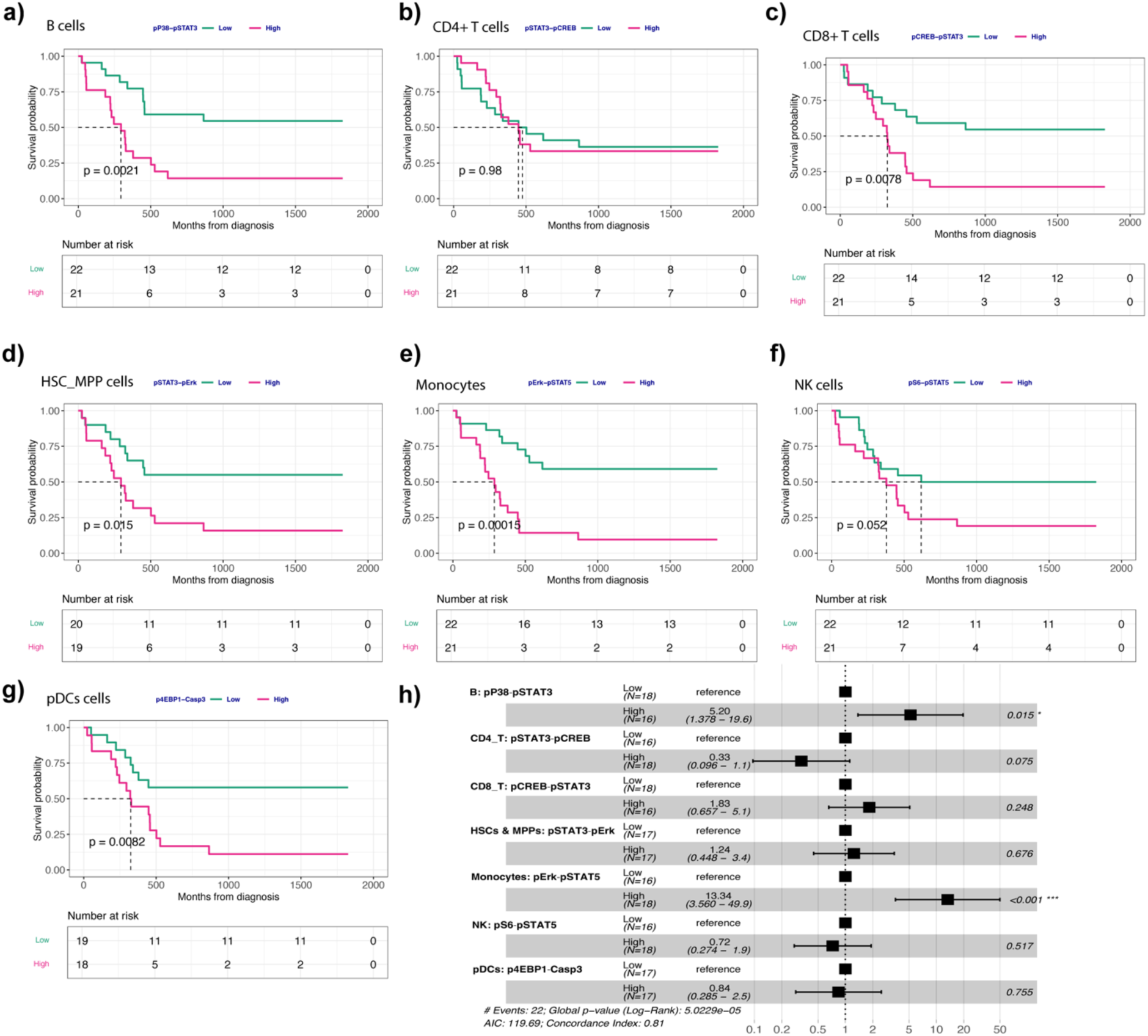
Independent survival analysis using the most important DREMI scores learnt from the XGBoost models. a) Survival probability of patients with pP38-pSTAT3 low vs high DREMI scores for B cells; b) Survival probability of patients with pSTAT3-pCREB low vs high DREMI scores for CD4_T cells; c) Survival probability of patients with pCREB-pSTAT3 low vs high DREMI scores for CD8_T cells; d) Survival probability of patients with pSTAT3-pErk pSTAT3 low vs high DREMI scores for HSC_MPP cells; e) Survival probability of patients with pErk-pSTAT5 low vs high DREMI scores for Monocytes; f) Survival probability of patients with pS6-pSTAT5 low vs high DREMI scores for NK cells; g) Survival probability of patients with p4EBP1-Casp3 low vs high DREMI scores for pDCs; h) Cox proportional hazards modelling using as input the most important cell type-specific DREMI scores learnt from the ML models. For subplots a-g univariate survival analysis is performed using Kaplan-Meier method.

**Supplementary Figure 15:**
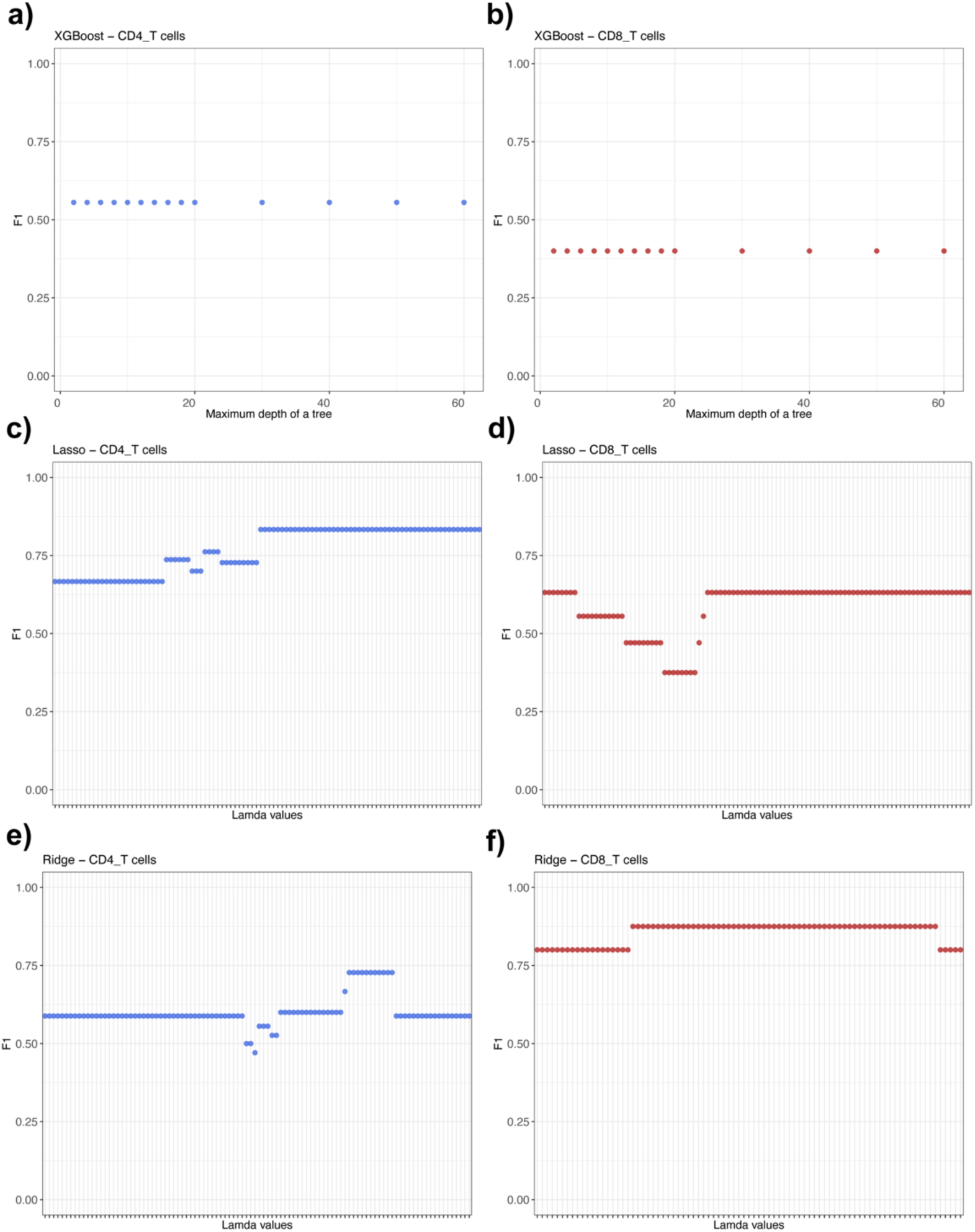
Optimising the internal parameters of the classification algorithms used for survival prediction. a-b) Snapshot of the optimisation process for XGBoost’s parameter maximum depth of trees; c-d) Snapshot of the optimisation process for LASSO’s parameter lamda; e-f) Snapshot of the optimisation process for Ridge regression’s parameter lamda. In all cases CD4_T and CD8_T cell type modelling is shown.

## Supplementary Tables

**Supplementary Table 1:** Characteristics of leukemia patients used in the case study.

| Patient ID | original patient nr from ref. 21 | Age | 5-year survival (days) |
| --- | --- | --- | --- |
| P1 | P1 | 35-60 | Alive |
| P2 | P2 | >60 | Alive |
| P3 | P3 | <35 | Alive |
| P4 | P4 | >60 | 456 |
| P5 | P5 | 35-60 | Alive |
| P6 | P6 | >60 | Alive |
| P7 | P7 | >60 | 246 |
| P8 | P8 | 35-60 | 189 |
| P9 | P9 | <35 | 286 |
| P10 | P10 | >60 | Alive |
| P11 | P11 | >60 | 328 |
| P12 | P12 | >60 | 502 |
| P13 | P13 | >60 | Alive |
| P14 | P14 | 35-60 | 448 |
| P15 | P15 | >60 | 378 |
| P16 | P16 | >60 | 221 |
| P17 | P17 | >60 | 56 |
| P18 | P18 | 35-60 | Alive |
| P19 | P19 | >60 | 51 |
| P20 | P20 | 35-60 | Alive |
| P21 | P21 | 35-60 | 24 |
| P22 | P22 | 35-60 | 324 |
| P23 | P23 | >60 | 447 |
| P24 | P25 | >60 | 229 |
| P25 | P26 | >60 | 162 |
| P26 | P27 | >60 | 321 |
| P27 | P28 | 35-60 | 48 |
| P28 | P29 | >60 | Alive |
| P29 | P30 | >60 | Alive |
| P30 | P31 | 35-60 | Alive |
| P31 | P33 | 35-60 | 457 |
| P32 | P34 | 35-60 | 528 |
| P33 | not included | >60 | 223 |
| P34 | not included | >60 | 295 |
| P35 | not included | 35-60 | 339 |
| P36 | not included | >60 | 187 |
| P37 | not included | >60 | 55 |
| P38 | not included | 35-60 | 26 |
| P39 | not included | 35-60 | Alive |
| P40 | not included | 35-60 | Alive |
| P41 | not included | 35-60 | Alive |
| P42 | not included | 35-60 | 618 |
| P43 | not included | 35-60 | 865 |

**Supplementary Table 2:**
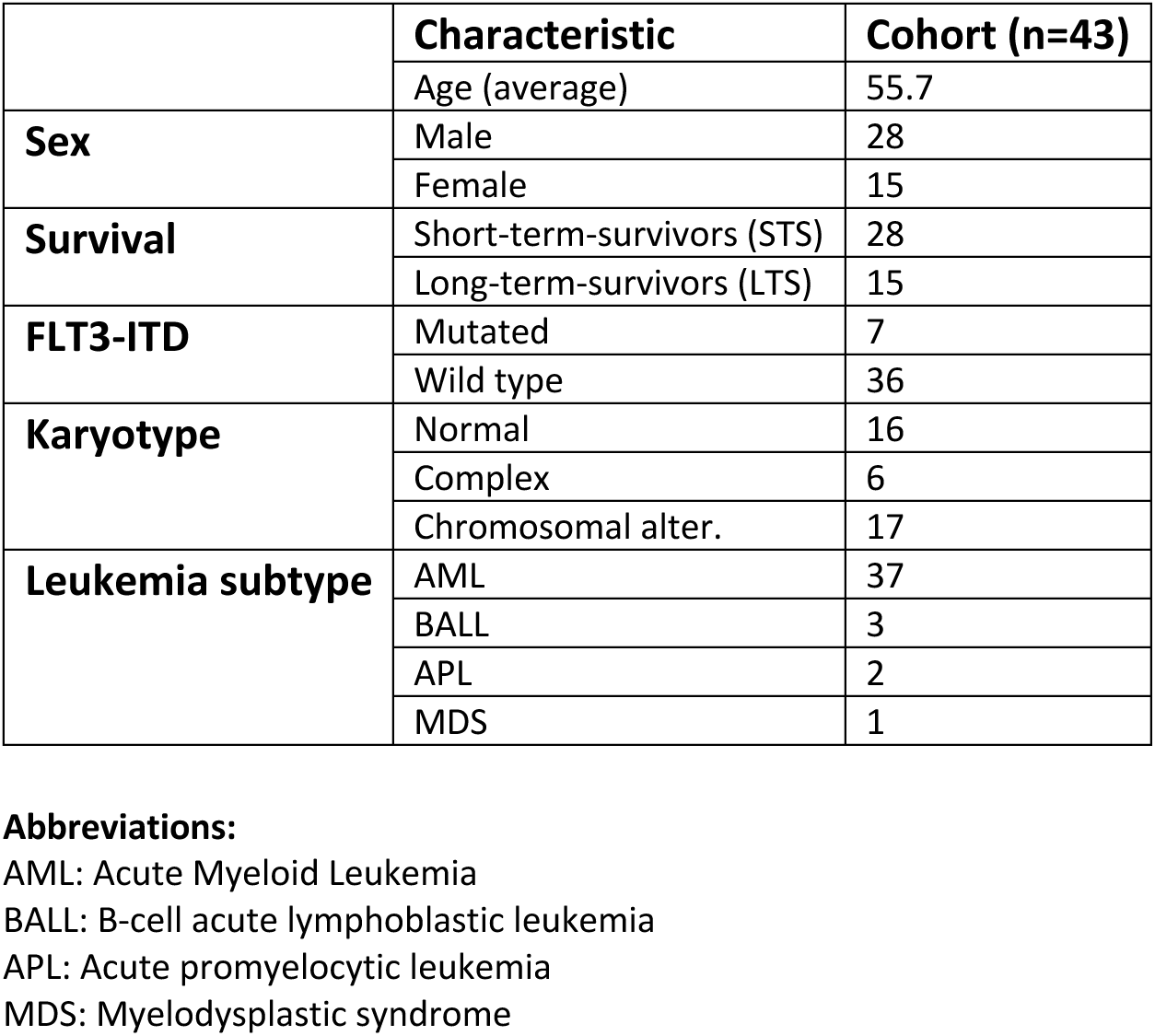
Cohort summary statistics.

